# Sustained Androgen Receptor signaling is a determinant of melanoma cell growth potential and tumorigenesis

**DOI:** 10.1101/2020.05.26.116665

**Authors:** Min Ma, Soumitra Ghosh, Daniele Tavernari, Atul Katarkar, Andrea Clocchiatti, Luigi Mazzeo, Anastasia Samarkina, Justine Epiney, Yi-Ru Yu, Ping-Chih Ho, Mitchell P. Levesque, Berna C. Özdemir, Giovanni Ciriello, Reinhard Dummer, G. Paolo Dotto

**Author notes:** Co-first authors (Min Ma, Soumitra Ghosh).

## Abstract

Melanoma is a benchmark of major clinical significance for cancer development with greater aggressiveness in the male than the female population. Surprisingly little is known on the role of androgen receptor (AR) signaling in the disease. Irrespectively of expression levels, genetic and pharmacological suppression of AR activity in a large panel of melanoma cells, derived from both male and female patients, suppresses proliferation and self-renewal potential while, conversely, increased AR expression or ligand stimulation enhance proliferation. AR gene silencing in multiple melanoma lines elicits a shared gene expression signature related to interferon- and inflammatory cytokines signaling with an inverse association with DNA repair-associated genes, which is significantly linked with better patients’ survival. AR plays an essential function in maintenance of genome integrity: in both cultured melanoma cells and tumors, loss of AR activity leads to chromosomal DNA breakage, leakage into the cytoplasm, and stimulator of interferon genes (STING) activation. *In vivo*, reduced tumorigenesis resulting from AR gene silencing or pharmacological inhibition is associated with intratumor macrophage infiltration and, in an immune competent mouse model, cytotoxic T cell activation. Although at different levels, androgens are produced in both male and female individuals and AR targeting provides an attractive therapy approach for improved management of melanoma irrespective of patients’ sex and gender.

**Significance:** The study uncovers an essential role of androgen receptor (AR) signaling in melanoma cell expansion and tumorigenesis, with loss of AR activity inducing cellular senescence, genomic DNA breakage, a STING dependent inflammatory cascade and immune cells recruitment. Use of AR inhibitors as growth inhibitory and DNA damaging agents in melanoma cells can provide an attractive venue for new combination approaches for management of the disease.

## Introduction

Malignant melanoma is the fifth-most common cancer in the world, and its incidence is rising. Among the many prognostic risk factors that have been proposed for the disease, one of the most intriguing and less understood is sex (1). In fact, melanoma is an example of primary clinical significance for investigating sex-related differences in cancer incidence and survival, with the male population having greater susceptibility than the female, across all ages (1). Although differences in lifestyle and behavior may explain the delay and higher disease stage in men at diagnosis, the female survival advantage persists even after adjusting for these and additional variables (histological subtypes, Breslow thickness, body site) (2, 3).

Differences in sex hormone levels and their downstream pathways play a key role in sexual dimorphism in multiple cancer types (4), including melanoma (1). As for sexual dimorphism in other cancer types (4), even for susceptibility to melanoma differences in sex hormone levels and/or downstream pathways are likely to play a key role (1). Relative to sex protein hormones, much more evidence exists on the impact of sex steroid hormones on cancer development (4). The great majority of accrued information for melanoma relates to estrogen signaling, while much less is known on androgen signaling.

In experimental settings, estrogen signaling was found to restrict melanocyte proliferation, enhance differentiation and suppress melanoma development (5–7). In spite of the experimental evidence, epidemiological studies on the interconnection between estrogen levels and melanoma development and progression yield conflicting conclusions (1) (7), which may be due, in part, to the difficulty in controlling for estrogen levels, which vary with the menstrual cycle, onset of menopause, use of oral contraceptives and hormone replacement therapy. Additionally, the possible interplay between estrogens and other hormones, specifically androgens, has not been taken into consideration. An interplay with frequently opposite effects between estrogen and androgen signaling has been reported for several cell types (4), which may extend to melanocytes.

The androgen receptor (AR) is expressed in many cell types and, while most studies have focused on prostate cancer, AR signaling has been implicated in tumorigenesis in other organs, specifically breast, bladder, kidney, lung, and liver (8). Surprisingly little is known on the role of androgen receptor signaling in melanoma. As early as 1980, it has been proposed that differences in androgen levels could explain the lower survival of male melanoma patients than females (9). Since then, however, only circumstantial pharmacological evidence has been obtained, pointing to a positive role of AR signaling in development of the disease (1). For instance, in a human melanoma cell line expressing an atypical form of AR, incubation with androgens significantly stimulated proliferation, with effects that were reversed by treatment with the androgen antagonist flutamide (or its active metabolite hydroxyflutamide) (10). The nonsteroidal antiandrogen flutamide was also found to be effective in diminishing tumor growth, and increasing survival of nude mice inoculated with human melanoma cells through possibly indirect effects (11). In fact, other work reported that administration of flutamide increased murine splenocyte proliferation, and interferon secretion, in response to irradiated murine B16 melanoma cells, and when flutamide was administered with an irradiated B16 vaccine, this combination improved the survival of mice implanted with non-irradiated B16 cells (12). Despite the above, genetic evidence in support of an intrinsic role of AR signaling in melanoma development is missing, with the possible exception of a study of a melanoma cell line plus/minus infection with a single shRNA silencing vector, which resulted in limited AR down-modulation (13). AR signaling in this setting was implicated in control of melanoma cells invasive properties, without any effect on proliferation.

In this study, based on analysis of a large panel of clinical samples and melanoma cells from both male and female patients, we show that, irrespective of expression levels, genetic and pharmacological suppression of AR activity triggers melanoma cell senescence and limits tumorigenesis, eliciting a gene expression signature related to interferon- and inflammatory cytokines and associated with better patients’ survival. Loss of AR activity in both melanoma cells and tumors is sufficient to cause massive chromosomal DNA breakage and leakage into the cytoplasm, with a stimulator of interferon genes (STING) dependent inflammatory signaling cascade, intratumor macrophage infiltration and, in an immune competent mouse model, cytotoxic T cell activation. As androgens are produced in both male and female individuals, AR represents an attractive target for improved management of the disease in patients of the two sexes.

## Results

### AR is heterogeneously expressed in melanocytic lesions and melanoma cells

Melanoma tumors are characterized by distinct phenotypic states and display significant intra and inter-tumor heterogeneity (14). Double immunofluorescence (IF) analysis of melanocytes in benign or dysplastic nevi or metastatic melanoma versus melanocytes from flanking skin showed consistently increased levels of AR expression in the melanocytic lesions, with heterogeneity of AR protein expression at the single-cell level (Figure 1A, Supplementary Figure 1). Further double IF analysis of these and additional individual lesions at three different positions within the lesions as well as melanoma tissue microarrays showed variable degrees of AR expression irrespectively of stages of neoplastic development, sex and age of patients (Figure 1B, C, Supplementary Figure 2, 3 and Supplementary Table 1). Immunohistochemical staining confirmed the IF results with prevalent nuclear AR localization in lesions with elevated and intermediate expression and more uneven localization when lowly expressed (Figure 1D, Supplementary Figure 4).

**Figure 1.**
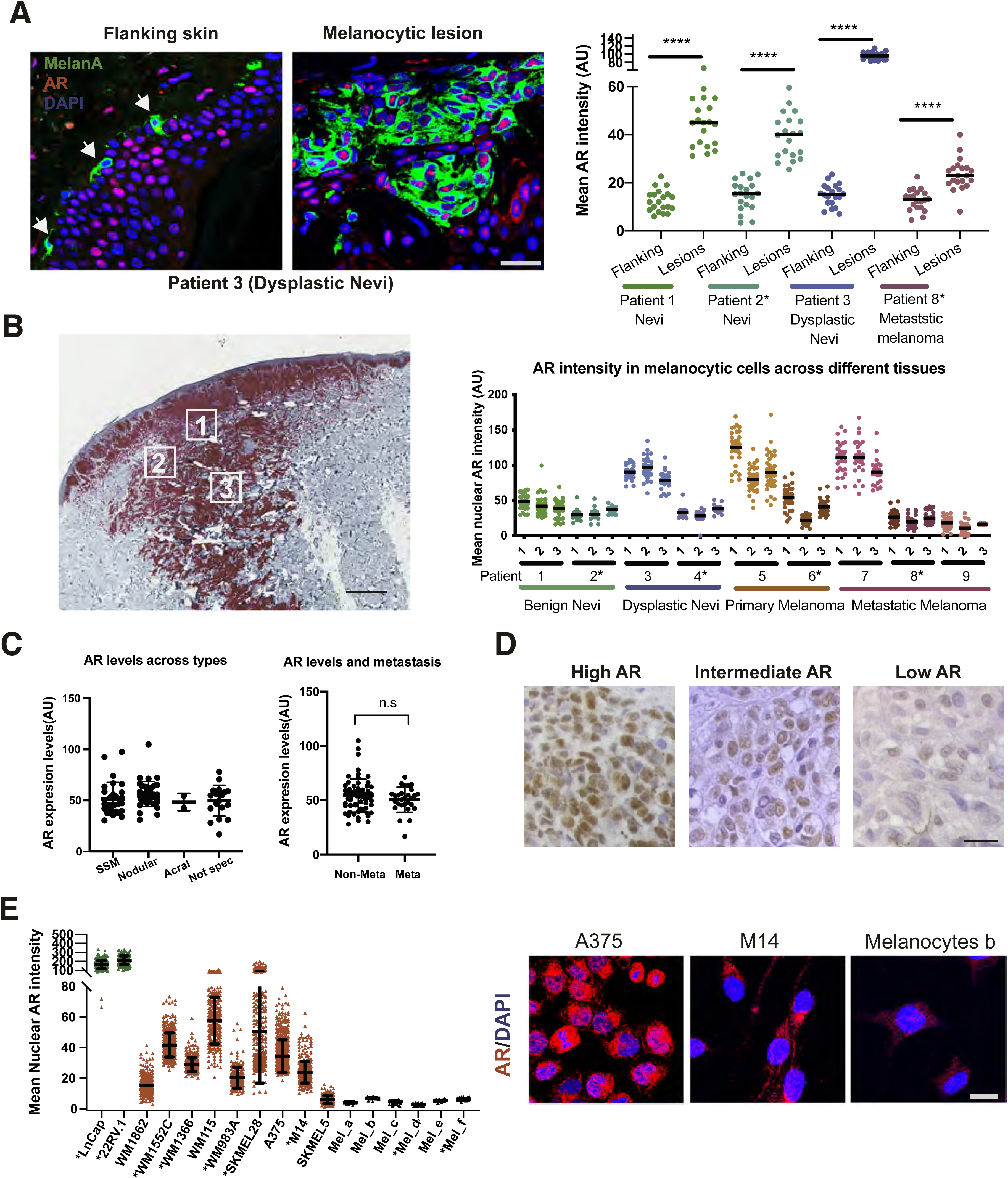
AR expression in melanoma cells. A) <u>Left</u>: Representative images of AR expression in cells of melanocytic lesions versus melanocytes in flanking normal skin (arrows) stained by double immunofluorescence with antibodies against AR (red) and MelanA (green) for melanocytes identification, with DAPI staining for nuclear localization (blue). Scale bar 30 µm. Additional images of this and other lesions are shown in Supplementary Figure 1. <u>Right</u>: Quantification of nuclear AR fluorescence signal in individual MelanA-positive cells (dots) of benign nevi, dysplastic nevi and metastatic melanomas versus melanocytes of flanking skin (samples from male patients in this and following panels are indicated by asterisks). Arbitrary fluorescence intensity values (AU) per individual cells are indicated together with mean and statistical significance (unpaired t test, **** < 0.001). B) <u>Left</u>: Immunohistochemical staining of a primary melanoma lesion with anti-MelanA antibodies and topographically distinct areas (boxes1, 2 and 3) analyzed for single cell AR expression by double immunofluorescence of parallel sections with anti AR and MelanA antibodies. Scale bar 500 µm. Immunofluorescence images of cells in this and other lesions are shown in Supplementary Figure 2. <u>Right</u>: Quantification of nuclear AR fluorescence signal in individual MelanA-positive cells (dots) from three topographically delimited areas per lesion (numbered as in B) of benign nevi, dysplastic nevi, primary and metastatic melanomas from different patients. Arbitrary fluorescence intensity values (AU) per individual cells are indicated together with the mean. C) Quantification of AR fluorescence signal in MelanA-positive cells in a tissue microarray of different types of melanoma patients with different clinical histories (left) and of metastatic and non-metastatic form (right). SSM = Superficial Spreading Melanoma. Acral = acral lentiginous melanoma. Quantification was based on digitally-acquired images of three independent fields per clinical lesion (a minimum of 50 cells per field) on the arrays. Results are expressed as average values for each lesion (dots) together with mean per type of lesions and primary versus metastatic. Quantification of samples divided by sex and age of patients is provided in Supplementary Figure 3. Patient sample details together with clinical diagnosis, age and sex is provided in Supplementary Table 1. D) Immunohistochemical staining with anti-AR antibodies of melanomas with high versus intermediate and low AR expression as assessed by double immunofluorescence analysis in (B). Scale bar: 30 µm. Lower magnification images with Melan A staining of parallel sections are shown in Supplementary Figure 4. E) <u>Left</u>: Quantification of nuclear AR expression by immunofluorescence analysis of the indicated melanoma cell lines or primary melanoma cells (red) versus primary melanocytes (black), and prostate cancer cell lines (green) examining > 100 cells per sample. Values for individual cells are indicated as dots together with mean ± SD. <u>Right</u>: Representative images of melanoma cells (A375, M14) and primary melanocytes with high versus low AR expression. Scale bar: 10 µm. Additional images of cells are shown in Supplementary Figure 5.

Immunostaining of cultured cells showed also a variation in AR protein expression among various melanoma cell lines and primary melanoma cells derived from male or female patients, with AR levels being uniformly low in primary melanocytes (Figure 1E, Supplementary Figure 5, Supplementary Table 2). As observed *in vivo*, AR localization was largely nuclear in melanoma cells with elevated expression, similar to LnCAP or 22RV.1 prostate cancer cells, while in melanoma cells or primary melanocytes with low AR levels, there was limited punctate nuclear localization with prevalent peri-nuclear distribution (Figure 1E, Supplementary Figure 5).

Variations in AR expression were further confirmed by immunoblotting with two different antibodies, with a similar pattern of bands, and by RT-qPCR analysis of melanoma cell lines, early passage primary melanoma cells and primary melanocytes, which were again found to be more lowly expressing (Supplementary Figure 6).

### Sustained AR expression is required for melanoma cell proliferation and self-renewal potential

The heterogeneous levels of AR expression raised the question of its biological significance. Accordingly, we silenced *AR* expression in a panel of melanoma cells harboring either *BRAF* or *NRAS* mutations individually and in combination with *TP53, PTEN* and/or *CDK4* mutations (Supplementary Figure 7). Irrespectively of basal levels of AR expression, silencing of the gene by two different shRNAs resulted in all cases in drastically reduced proliferation and self-renewal as assessed by cell density, clonogenicity and sphere formation assays (Figure 2A-C, Supplementary Figure 8). Effects were paralleled by decreased DNA synthesis, induction of apoptosis and cellular senescence (Figure 2D-F, Supplementary Figure 9). The shRNA gene silencing effects were suppressed in melanoma cells concomitantly infected with an AR over-expressing lentivirus (Figure 2G, Supplementary Figure 10), which was by itself sufficient to enhance proliferation of primary melanocytes as well as melanoma cells with low AR expression (Figure 2H).

**Figure 2.**
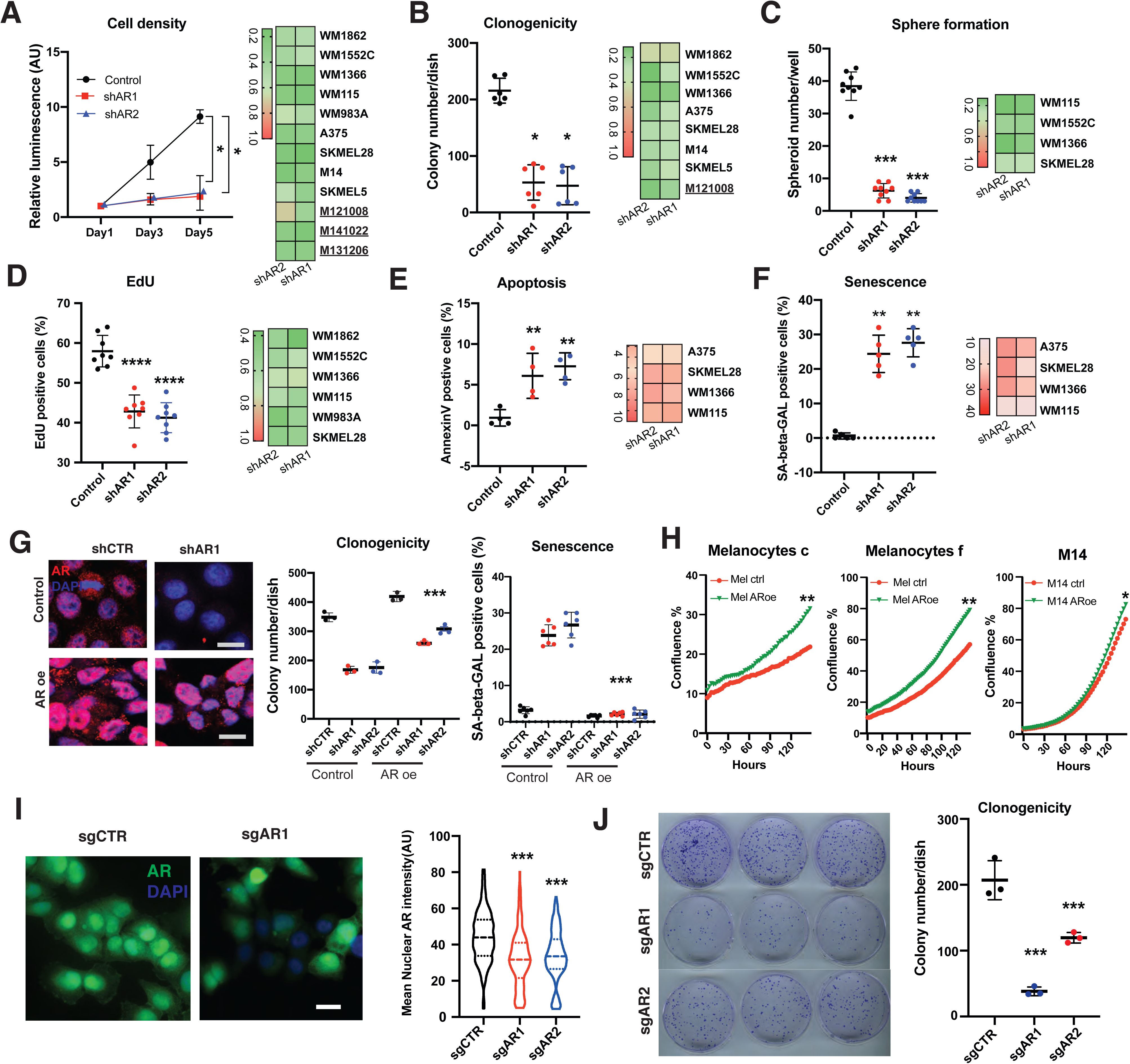
AR down-modulation suppresses melanoma cell proliferation and self-renewal potential. A) <u>Left</u>: WM1366 melanoma cells stably infected with 2 AR-silencing lentiviruses versus empty vector control were analyzed by cell density assays (CellTiter-Glo) at the indicated days after infection and selection. Results are presented as luminescence intensity values relative to day 1. Data are shown as mean ± SD, 1-way ANOVA with Dunnett’s test. n = 3 independent experiments. *P < 0.05. <u>Right</u>: results with additional melanoma cell lines and primary melanoma cells (<u>MM141022</u>, <u>M131206</u>, <u>M121008</u>) presented as heat maps. Efficiency of AR gene silencing is shown in Supplementary Figure 7. Corresponding individual plot results together with statistical significance are shown in Supplementary Figure 8A. B, C) Proliferation potential of the indicated melanoma cells plus/minus AR silencing was assessed, at day 6 after lentivirus infection, by clonogenicity (B) or sphere formation assays (C). For each condition, cells were tested in triplicate dishes, with all experiments repeated 3 times. Results for WM1366 melanoma cells are shown as individual cultured dishes together with mean of ± SD, 1-way ANOVA with Dunnett’s test. n = 3 technical replicates (experiments). *P < 0.05; ***P < 0.005. Results for other melanoma cells are presented as heat map, with individual plots for each cell line and representative images being shown in Supplementary Figure 8B, C. D-F) Melanoma cells plus/minus AR silencing as in the previous panels were tested by EdU labelling assay (D), apoptosis by annexin V staining (E) or senescence associated ß-Gal activity (F). <u>Left</u>: quantification of the percentage of positive cells in individual cultured dishes of WM1366 melanoma cells plus/minus shAR together with mean of ± SD, 1-way ANOVA with Dunnett’s test. n = 3 independent experiments). ** P < 0.01; ****P < 0.001. <u>Right</u>: results with additional melanoma cell lines presented as heat maps with individual plots for each cell line shown in Supplementary Figure 9. G) <u>Right</u>: representative IF analysis of AR expression in A375 cells stably infected with an AR-over-expressing lentivirus or vector control and super-infected with an AR silencing lentivirus or corresponding control. Scale bar 10 µm. Quantification of results, also in cells infected with a second AR silencing lentivirus, together with mRNA expression measurements are shown in Supplementary Figure 10A and B. <u>Left</u>: clonogenicity and senescence associated ß-Gal activity assays of A375 melanoma cells plus/minus AR silencing and overexpression as indicated. Data are shown as mean ± SD, 1-way ANOVA with Dunnett’s test. n = 3 dishes, *** p<0.005. Cell density, EdU labelling and apoptosis assays with A375 cells plus/minus concomitant AR overexpression and silencing are shown in Supplementary Figure 10C-E. H) Proliferation live-cell imaging assays (IncuCyteTM system, Essen Instruments) of the indicated primary melanocyte strains (c, f) and melanoma cells (M14) stably infected with an AR over-expressing lentivirus versus empty vector control. Cells were plated in triplicate wells in 96-well plates followed by cell density measurements (4 images per well every 4 hours for 128 hours). n (number of wells) = 3, Pearson r correlation test. *P < 0.05; ** P < 0.01. I) Representative images and quantification of AR expression by IF analysis of dCas9-KRAB-expressing A375 cells infected with lentiviruses expressing two single guide RNAs targeting the AR promoter region (sgAR1, sgAR2) versus scrambled single guide RNA control (sgCTR) for 3 day. Scale bar 10 µm. Violin plot showed the nuclear AR fluorescence intensity, together with mean ± SD, n > 300 cells per sample, 1-way ANOVA with Dunnett’s test. *** p<0.005. J) Parallel cultures of cells as in the previous panel were tested by clonogenicity assays on triplicate dishes, starting at day 3 after single guide RNA expression. n = 3 biological replicates (dishes), 1-way ANOVA with Dunnett’s test. *** p<0.005.

As an alternative to shRNA-mediated gene silencing, we also downmodulated AR expression by a CRISPRi system (15, 16), whereby a dCas9-KRAB transcription repressor was directed to the AR promoter region by two different guide RNAs (sgRNAs). Mass infection of dCas9-KRAB expressing melanoma cells with two lentiviruses with AR-targeting sgRNAs reduced significantly AR protein levels and decreased clonogenicity, reproducing the effects of AR gene silencing (Figure 2I, J).

### Modulation of melanoma and melanocyte proliferation by pharmacological inhibition and agonist stimulation

AR is a fundamental target for therapy of metastatic prostate cancer, and inhibitors with multiple mechanisms of action and efficacy have been developed (17). Treatment of different melanoma cell lines with several AR inhibitors, including one that functions through both ligand-competitive and non-competitive mechanisms, AZD3514 (18), and another, pure ligand competitive inhibitor, enzalutamide (19), exerted similar growth suppressive effects although at different doses (Figure 3A). The first compound exhibited a greater potency, which we found to be associated, as previously reported for LNCaP cells (18), with down-modulation of AR expression in two of three tested cell lines (Supplementary Figure 11A). The AZD3514 inhibitory effects were confirmed by treatment of a larger panel of melanoma cell lines and primary melanoma cells with different levels AR expression, consistent with the basal protective function investigated below (Figure 3B, C, Supplementary Figure 11B, C).

**Figure 3.**
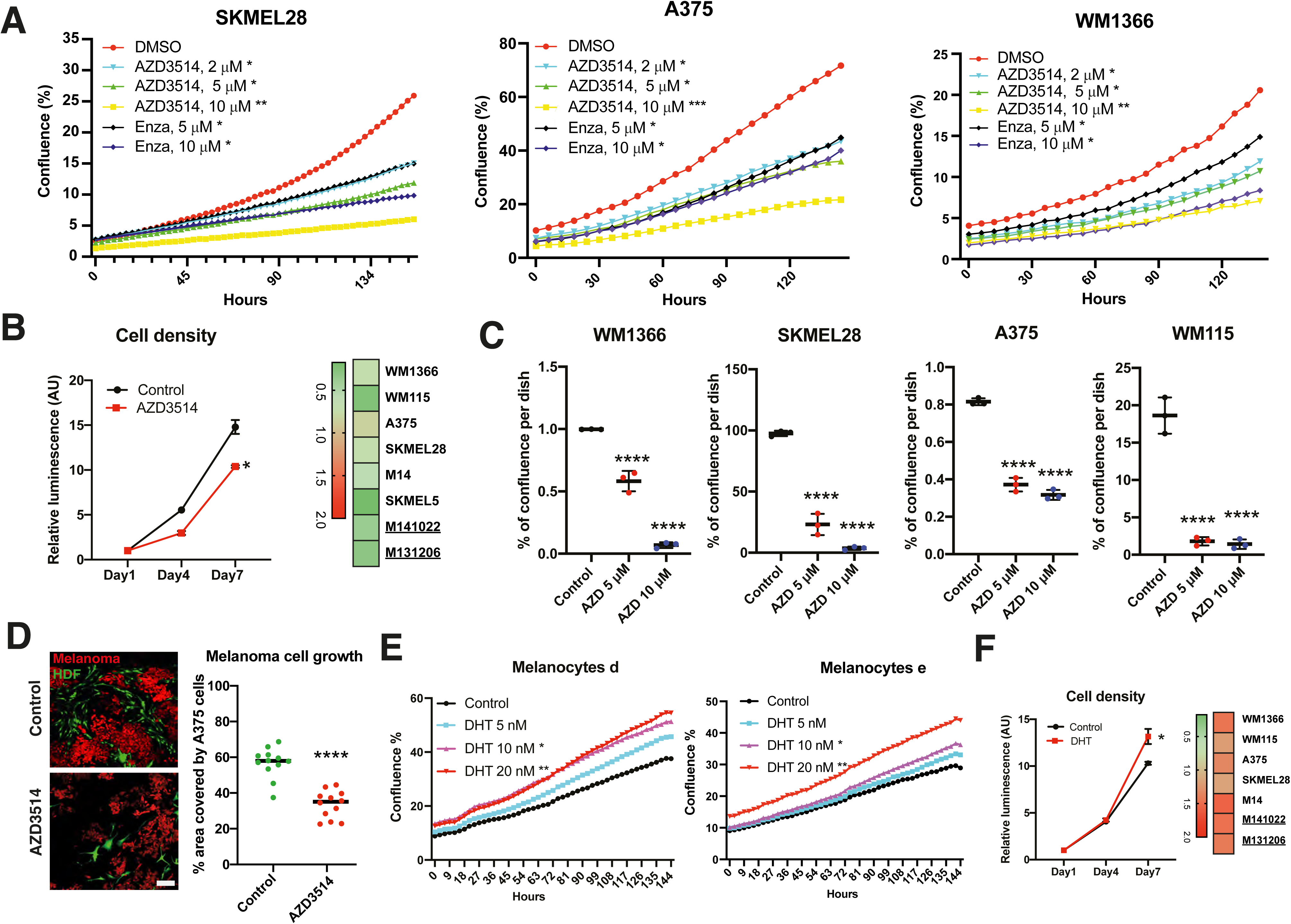
Modulation of melanoma cells proliferation by pharmacological inhibition and agonist stimulation. A) Proliferation live-cell imaging assays of the indicated melanoma cell lines treated with the AR inhbitors AZD3514 (2, 5, 10 µM) or Enzalutamide (5, 10 µM) versus DMSO control. Cells were plated on triplicate wells in 96-well dishes followed by cell imaging measurements (IncuCyteTM system, Essen Instruments), capturing 4 images per well every 4 hours for 140 hours). n (number of wells) = 3, Pearson r correlation test. *P < 0.05; ** P < 0.01; ***P < 0.005. B) <u>Left</u>: WM1366 melanoma cells treated with AZD3514 versus DMSO control were analyzed by cell density assays (CellTiter-Glo) at the indicated days. Results are presented as luminescence intensity values relative to day 1. Data are shown as mean ± SD, 1-way ANOVA with Dunnett’s test. n = 3 independent experiments. *P < 0.05. <u>Right</u>: results with additional melanoma cell lines and primary melanoma cells (MM141022, M131206) presented as heat maps, with individual plots per cell line being shown in Supplementary Figure 11B. C) The indicated melanoma cells in triplicate dishes were treated with AZD3514 (5 µM and 10 µM) versus vehicle control (DMSO) followed by crystal violent staining 7 days later and ImageJ determination of % of cell coverage area. Data are shown as mean ± SD, 1-way ANOVA with Dunnett’s test. n = 3 biological replicates (dishes). ****P < 0.001. D) *In vitro* cancer / stromal cell expansion assays, based on the co-culture of RFP-expressing A375 melanoma cells together with GPF-expressing human dermal fibroblasts (HDF) in Matrigel in triplicate dishes, plus treatment with AZD3514 (10 µM) or DMSO control for 4 days. Shown are representative images and quantification of melanoma cells expansion (percentage area covered by melanoma cells per field). Each dot represents one analyzed field. n (number of fields) = 12, two-tailed paired t test, ****P < 0.001. Scale bar: 30 µm. E) Proliferation live-cell imaging assays of two primary melanocyte strains cultured in medium with charcoal-stripped serum and treated with dihydrotestosterone (DHT) at the indicated concentrations versus DMSO control. Cells were plated in triplicate wells in 96-well plates and imaged using IncuCyteTM system (Essen Instruments), every 4 hours for 145 hours. n (number of wells) = 3, Pearson r correlation test. *P < 0.05; ** P < 0.01. Results of a similar assay with another primary melanocyte strain and melanoma cells are shown in Supplementary Figure 12A. F) <u>Left</u>: cell density assays (CellTiter-Glo) of WM1366 melanoma cells cultured in medium with charcoal-treated serum and treated with DHT (20 nM) versus DMSO control for the indicated days. Results are presented as luminescence intensity values relative to day 1. Data are shown as mean ± SD, 1-way ANOVA with Dunnett’s test. n = 3 independent experiments. *P < 0.05. <u>Right</u>: results of a similar assay with additional melanoma cell lines and primary melanoma cells (MM141022, M131206) presented as heat maps, with individual plots and statistical significance shown in Supplementary Figure 12B.

We recently reported that suppression of AR activity in human dermal fibroblasts (HDFs) by a ligand-competitive inhibitor induces expression of a battery of tumor promoting CAF effector genes, similarly to silencing of the gene (20). To assess the net effects of AR inhibitors on melanoma cells in the presence of surrounding HDFs, we resorted to an *in vitro* cancer / stromal cell expansion assay based on the co-culture in matrigel of fluorescently labelled cells (20). As shown in Figure 3D, expansion of melanoma cells admixed with HDFs was significantly reduced by treatment with the AR inhibitor AZD3514, consistent with the efficacy of this compound in the *in vivo* assays shown further below.

Conversely to the growth suppressing effects of the AR inhibitors, proliferation of primary melanocytes and melanoma cells in charcoal-stripped medium was significantly enhanced by treatment with the AR agonist dihydrotestosterone (DHT) in a dose-dependent manner (Figure 3E, Supplementary Figure 12A). Proliferation of other melanoma cell lines and primary melanoma cells in charcoal-stripped medium was also enhanced by DHT stimulation (Figure 3F, Supplementary Figure 12B,C) and, when cultured under very sparse conditions, their expansion was very highly dependent on the hormone (Supplementary Figure 12D). Thus, besides being required, increased AR signaling is a positive determinant of melanoma cells proliferation.

### AR is a master regulator of melanoma gene expression

AR controls transcription through both direct and indirect DNA binding mechanisms (21), which could affect intrinsic control of proliferative potential and survival of melanoma cells as well as their capability to modulate the tumor microenvironment. By transcriptomic analysis of three different melanoma lines, two with *BRAF* and one with *NRAS* mutations, we identified 155 genes that were significantly and concordantly modulated by *AR* silencing in all three (Figure 4A, Supplementary Table 3). The two most down-modulated genes were *CDCA7L*, coding for a transcriptional repressor and c-MYC interacting protein with shared oncogenic function (22, 23), and *SENP3*, coding for a protease with a key role in SUMO activation and removal, which affects activity of many DNA repair proteins as well as NF-κB- and STING-dependent transcriptional regulation of inflammatory genes (24, 25). Top up-regulated genes included some encoding proteins with key immunomodulatory functions, *ICAM1* (26) and *TLR4* and *6* (27), RNA helicases involved in interferon signaling cascades, *DDX58* (RIG-1) and *IFIH1* (Melanoma differentiation-associated factor 5) (28), and axonal-guiding secreted proteins with an extended role in immune regulation and angiogenesis, *SEMA3A* (Semaphorin 3A) (29) and *NTN4* (Netrin 4) (30) (Figure 4A, Supplementary Table 3). Genes encoding for transcriptional regulators of inflammatory signaling cascades, such as *NFKBIA*, *NFKB2*, *IRF9*, *STAT3* and *PARP9*, were also consistently up-regulated (Supplementary Table S3).

**Figure 4.**
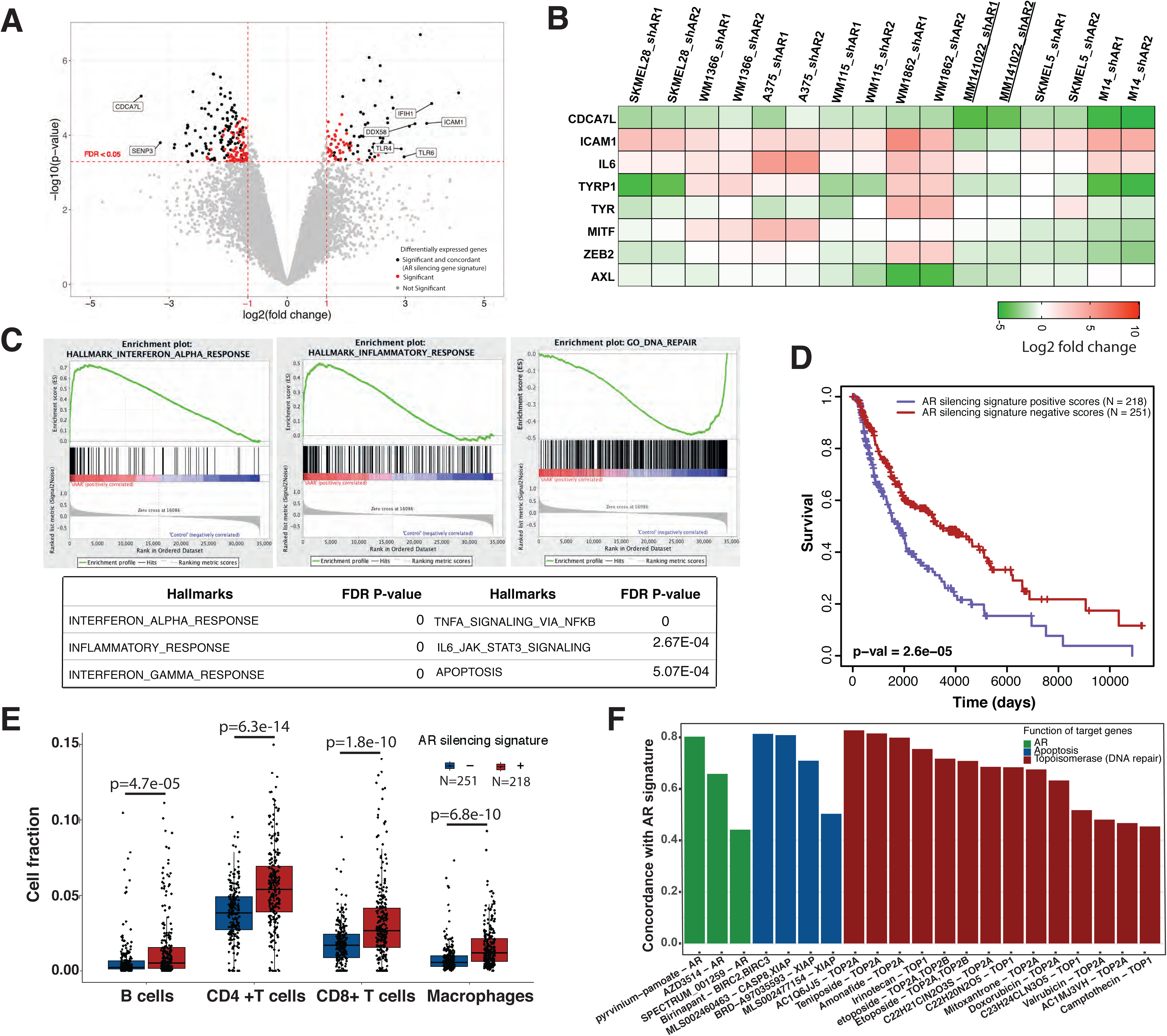
Global analysis of AR-regulated genes in melanoma cells and clinical relevance. A) Volcano plot showing the shared transcriptional changes elicited by AR silencing in WM1366, SKMEL28 and WM115 melanoma cells plus/minus AR-silencing by two different lentiviruses versus empty vector control. Cells were analyzed 5 days after infection by ClariomTM D array hybridization. The x-axis shows the log2 fold-change between the two conditions, the y-axis shows the –log10 (p-value). A False Discovery Rate (FDR) threshold of 0.05 and fold change thresholds of −1 and 1 are indicated by dashed red lines. Each dot represents one gene. Grey and red dots correspond to genes not significantly or non-concordantly modulated in the three melanoma lines, respectively. Black dots show genes above thresholds that are concordantly up- or down-regulated in all three cell lines and compose the AR silencing gene signature utilized for further analysis. A few selected genes among the most significantly differentially expressed ones are indicated. The list of 155 genes associated with AR silencing gene signature is provided in Supplementary Table 3. B) Expression of the indicated genes in multiple melanoma cell lines plus/minus AR silencing by two different lentiviruses versus empty vector control. RT-qPCR results after RPLP0 normalization are shown as a heat map of ratios of gene expression (folds of down- or up-regulation in green and red, respectively). Individual gene expression profiles (with most analyses based on two independent experiments) are shown in Supplementary Fig. S13. C) Gene set enrichment analysis (GSEA) of the common gene expression profiles elicited by AR gene silencing in WM1366, SKMEL28 and WM115 melanoma cells (Supplementary Table 3) versus a predefined set of gene signatures related to various processes and signaling pathways (Broad Institute, http://software.broadinstitute.org/gsea/msigdb/collections.jsp#H). Top: GSEA plot distribution of gene signatures related to interferon alpha, inflammatory response and DNA repair pathways. Genes are ranked by signal-to-noise ratio based on their differential expression in AR-silenced versus control melanoma cells; position of genes in the respective hallmark gene sets is indicated by black vertical bars, and the enrichment score is shown in green. Bottom: relevant gene sets most significantly associated with AR silencing gene signature are indicated together with the corresponding FDR-q values. The full list of significantly associated gene signatures is provided in Supplementary Table 4. D) Association of the AR silencing gene signature in melanoma cells - as obtained in panel A - with patients’ survival in TCGA Skin Cutaneous Melanoma (SKCM) dataset. Signature scores for each patient were computed from RNA-seq data with GSVA R package (40), with Kaplan-Meier curves showing that melanomas with positive scores for the AR silencing signature (red, N = 251) are significantly associated with better survival than the ones with negative scores, (blue, N = 218), p-value = 2.6e-05, log-rank test. E) Fraction of tumor-infiltrating immune cells estimated by EPIC R package analysis of TCGA Skin Cutaneous Melanoma (SKCM) dataset, using default reference profile in tumors positive and negative for the AR silencing signature (red and blue boxplots respectively). Cell fractions for B cells, CD4+ T cells, CD8+ T cells and macrophages are reported (each dot representing one tumor). Outliers with cell fraction greater than 0.15 are not shown. EPIC association plots for all other cell types are shown in Supplementary Figure 14A together with the enrichment scores of signature matrix associated with 22 different immune cell types determined by CIBERSORTx (Supplementary Figure 14B). F) Bar plot reporting the concordance between the melanoma AR silencing gene signature and iLINCS expression profiles of A375 cells treated with compounds targeting AR (green), apoptosis-related proteins (BIRC2, BIRC3, XIAP, blue) and topoisomerases (TOP1, TOP2A, red). Perturbagens of each class are sorted by concordance (p < 0.0001) and the names of the chemical compounds are reported on the x axis along with their molecular targets. If multiple signatures were available for the same perturbagen (e.g. by varying concentration or time), only the signature with the highest concordance was shown. A list of compounds eliciting gene expression profiles with concordance coefficient > 0.6 with AR silencing signature is reported in Supplementary Table 5.

The analysis was extended to a panel of other melanoma cell lines and primary melanoma cells with different levels of *AR* expression by RT-qPCR. *CDCA7L* expression was down-modulated by *AR* silencing in all tested cells, while ICAM1 was consistently up-regulated in parallel with IL6, a potent pro-inflammatory cytokine (Figure 4B, Supplementary Figure 13). As for "canonical" genes involved in melanoma progression, differentiation marker genes such as *TyR* and *TYRP1* were either up- or down-modulated by *AR* silencing in the various cell lines and so were the *MITF* master regulatory gene (31) and *ZEB2*, coding for a transcription factor with a role in melanogenesis upstream of MITF expression (32)*. AXL*, coding for a receptor tyrosine kinase implicated in melanoma aggressive behavior (33) was mostly down-modulated (Figure 4B, Supplementary Figure 13).

### The melanoma AR-dependent gene signature is of clinical relevance

By gene set enrichment analysis (GSEA) (34), gene signatures related to interferon- and inflammatory cytokines signaling as well as apoptosis were the most significantly associated with the common gene expression profile of melanoma cells with silenced *AR*, while an inverse association with DNA repair-associated gene signatures was noted (Figure 4C, Supplementary Table 4). Next, we defined an AR silencing gene signature comprising the 155 genes consistently modulated by AR loss in melanoma cells (Figure 4A, Supplementary Table 3) and assessed its clinical relevance by computing signature scores for 469 cutaneous melanoma patients in The Cancer Genome Atlas dataset (TCGA-SKCM). Patients were stratified as having positive scores (i.e. expression profiles similar to AR-silenced cell lines) or negative scores (i.e. expression profiles similar to AR-expressing cell lines). Patient with positive scores were found to have a significantly higher survival than those with negative scores across sexes (log-rank test, p-value = 2.6e-05; Figure 4D). The findings remained significant after correcting for age, sex, genomic subtype and primary or metastatic status (multivariate Cox regression, p-value = 0.002). Using the EPIC algorithm (35), we estimated the proportion of cancer and immune cells in the TCGA-SKCM melanoma samples cohort. We found a significantly higher proportion of B cells, CD4+ and CD8+ T cells and macrophages infiltrating melanomas with positive scores for the *AR* silencing signature than in melanomas with negatives scores (Figure 4E, Supplementary Figure14A). In order to validate and refine the analysis, we used an independent approach, CIBERSORTx (36), and estimated the proportion of 22 immune cell subtypes. With this approach, we found that tumors with positive *AR* silencing scores were selectively enriched for M1-like versus M2-like macrophages, and for CD4+ memory T cells (Supplementary Figure 14B).

The iLINCS (Integrative LINCS; http://www.ilincs.org/ilincs/) portal allows comparative analysis of transcriptional profiles of various cell lines in response to different drugs. A significant concordance was found between the *AR* silencing gene signature and the iLINCS-derived transcriptional profiles of A375 melanoma cells treated with several AR inhibitors as well as compounds inhibiting the anti-apoptotic BIRC2/3 and XIAP proteins and a number of DNA damaging agents targeting the TOPO2 and TOPO 1 enzymes (Figure 4F, Supplementary Table 5).

### AR loss triggers genomic DNA breakage, cytoplasmic leakage and STING-dependent gene expression

The "Stimulator of Interferon Genes" protein (STING) is a cytosolic DNA sensor with an important role in innate immunity (37). Its activation by the release of chromatin DNA fragments into the cytoplasm is a potent trigger of the interferon / inflammatory gene expression programs that we found to be induced by *AR* silencing in melanoma cells (38). Comet assays showed a striking induction of chromosomal DNA breakage by *AR* gene silencing in several melanoma cells, irrespective of their high or low levels of AR expression (Figure 5A), which was accompanied by increased levels of γ-H2AX, a marker of the DNA damage response (39) (Figure 5B, C). In parallel, *AR* silencing resulted in the abundant release of dsDNA fragments into the cytoplasm together with the induction of STING protein expression and aggregation, two signs of STING activation, as well as upregulation of the downstream immune-modulatory gene product ICAM1 (37) (Figure 5B, C). Similar observations were obtained upon treatment of melanoma cells with the AR inhibitor AZD3514 (Figure 5D, E). The findings are of functional significance as induction of AR target genes with key immune-modulatory functions, such as IL6 and ICAM1, was suppressed at both protein and mRNA levels by concomitant *AR* and *STING* knock-down (Figure 6A-C). The link between AR loss and ensuing events was further supported by the fact that chromosomal DNA damage and leakage into the cytoplasm, STING activation and IL6 and ICAM1 induction were all suppressed in cells in which AR gene silencing was counteracted by over-expression (Figure 6D, E).

**Figure 5.**
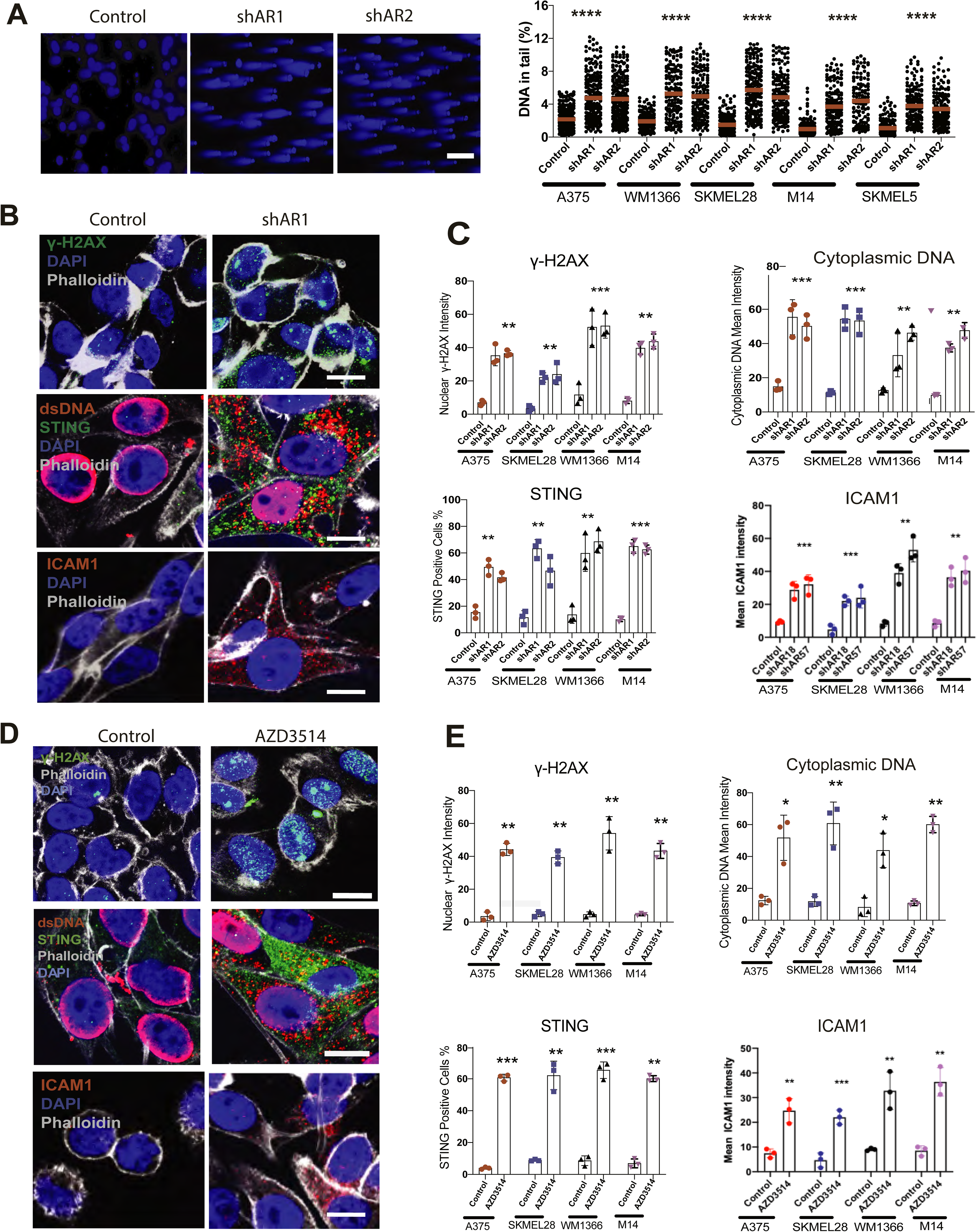
Loss of AR function induces DNA breakage, cytoplasmic dsDNA leakage and STING activation. A) Comet assays of multiple melanoma cell lines plus/minus shRNA-mediated AR silencing (5 days after infection). Shown are representative images of WM1366 melanoma cells together with quantification of % tail DNA (calculated by Comet Score 1.6.1.13 software, www.rexhoover.com) in five different melanoma cell lines. Scale bar: 10 µm. n (number of cells) =125; one-way ANOVA; ****p < 0.001. B, C) Double immunofluorescence image analysis of a panel of melanoma cell lines plus/minus shRNA-mediated AR silencing (5 days after infection). B) representative images of WM1366 stained with antibodies against γ-H2AX (green) and Phalloidin (gray) for cell border identification (upper panel), and dsDNA (red) and STING (green) (middle panel) and ICAM1 (red) (lower panels). Scale bar: 10 µm. C) quantification of nuclear γ-H2AX, cytoplasmic DNA, ICAM1 immunofluorescence signal intensity and percentage of STING positive cells in the indicated panel of melanoma cell lines plus/minus AR silencing. > 100 cells were counted in each condition. Results are expressed as mean ± SD. n (number of experiments) = 3; 1-way ANOVA with Dunnett’s test, ** P < 0.01, *** P < 0.005. D, E) Double immunofluorescence image analysis of a panel of melanoma cells treated with AZD3514 (10 µM) versus DMSO control for 2 days. Shown are representative images (D) and quantification (E) of the results as in the previous panel. n (number of experiments) = 3; two-tailed paired t-test, *P < 0.05; ** P < 0.01, *** P < 0.005.

**Figure 6.**
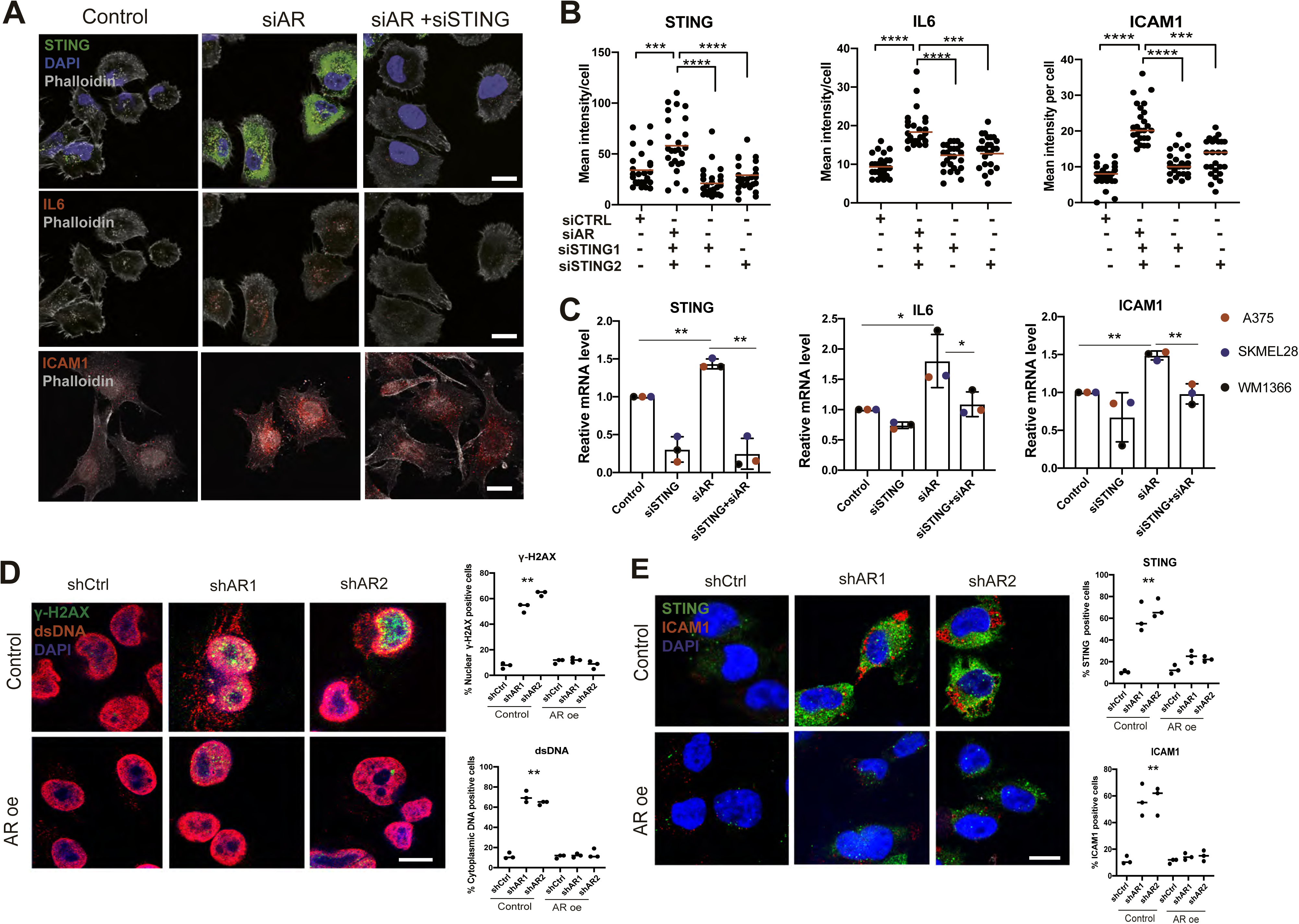
Loss of AR function induces STING-dependent gene expression. A, B) Counteracting impact of AR and STING gene silencing on IL6 and ICAM1 expression. Double immunofluorescence analysis of WM1366 melanoma cells transfected with STING and/or AR silencing siRNAs versus scrambled controls, with antibodies against STING (upper panel, green), IL6 and ICAM1 (middle and lower panels, red) with phalloidin staining for cell border delimitation (gray). Shown are representative images (A) and quantification (B) of STING, IL6 and ICAM1 fluorescence signal intensity per cell, 48 h after transfection. Each dot corresponds to mean fluorescence intensity per cell. n (number of cells) = 25; paired t-test, ***P < 0.005, ****P < 0.001. Scale bar: 10 µm. C) RT-qPCR analysis of STING, IL6 and ICAM1 mRNA expression in the indicated melanoma cell lines 48 h after transfection with STING and/or AR silencing siRNAs versus scrambled controls. Each bar corresponds to mean expression levels per melanoma cell line. Data are represented as mean ± SD. n (number of strains) = 3; 1-way ANOVA with Dunnett’s test, *P < 0.05, **P < 0.01. D) Representative double IF images and quantification of γ-H2AX expression (green) and cytoplasmic dsDNA (red) leakage in A375 cells stably infected with an AR-overexpressing (AR oe) or control lentivirus and super-infected with two AR silencing lentiviruses versus control. Scale bar: 10 µm. Data are from triplicate experiments; each dot represents one experiment. n (number of experiments) = 3; 1-way ANOVA with Dunnett’s test, ** P < 0.01. E) representative double IF images and quantification of STING (green) and ICAM1 (red) expression in A375 cells plus/minus AR overexpression and silencing as in the previous panel. n (independent experiments) = 3; 1-way ANOVA with Dunnett’s test, ** P < 0.01.

### AR loss or inhibition results in reduced tumorigenicity with enhanced inflammatory infiltrations

To assess the *in vivo* significance of the findings, we resorted to an orthotopic model of melanoma formation based on the intradermal injection of melanoma cells embedded in matrigel (20), which enables the assessment of early steps of tumor formation and expansion. Utilizing this assay, we found that tumorigenic expansion and proliferative activity of multiple melanoma cell lines (WM1366, A375 and SKMEL28) was significantly reduced, in male and female mice, by *AR* silencing (Figure 7A, B, Supplementary Figure 16, 17). In parallel, *AR* silencing resulted in cytoplasmic dsDNA release, STING aggregation and ICAM1 induction (Figure 7C-E). While host macrophages were mostly excluded by tumors formed by control cells, they actively infiltrated tumors formed by cells with silenced *AR* (Figure 7F; Supplementary Figure 15).

**Figure 7.**
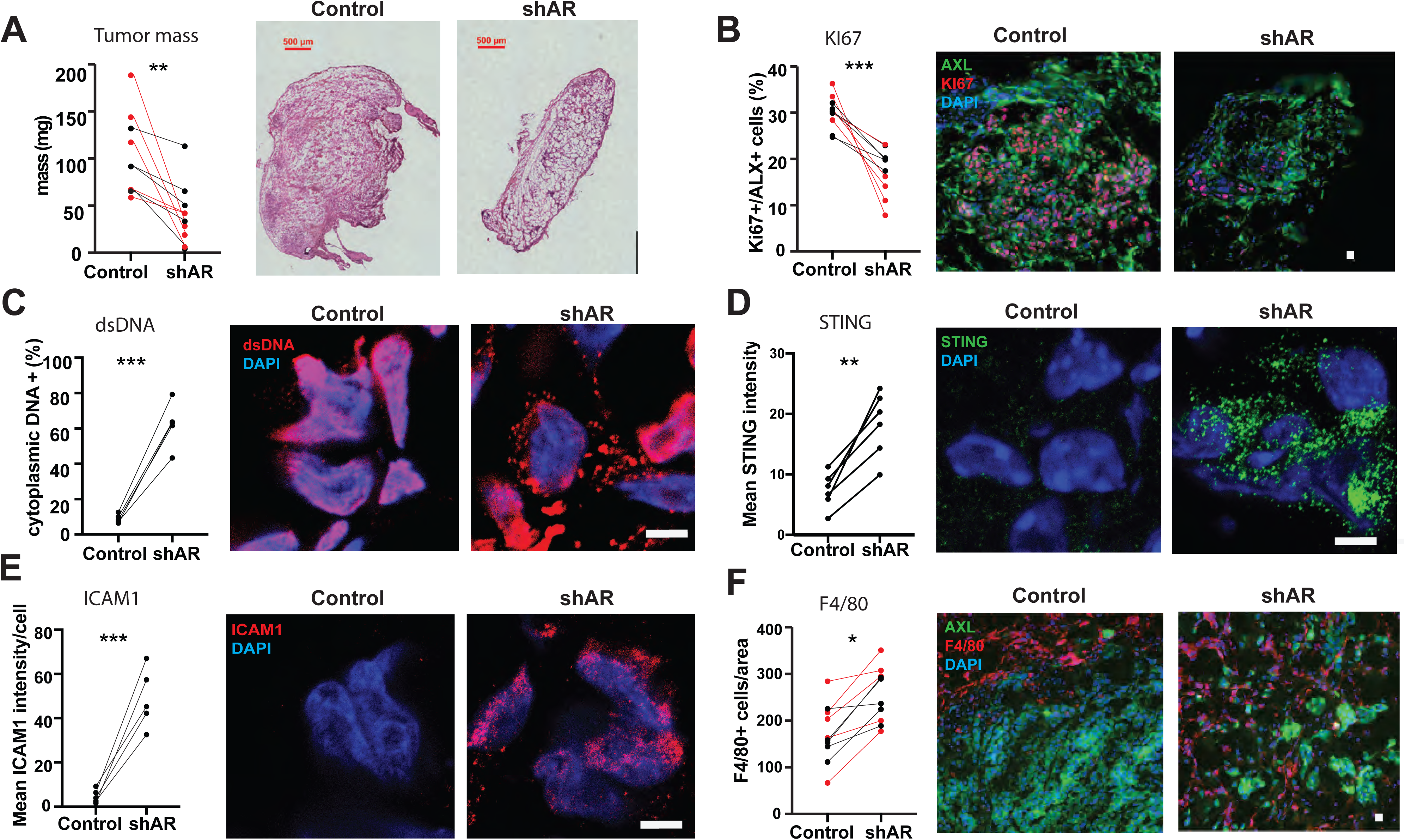
Suppression of melanoma formation by AR silencing. WM1366 melanoma cells infected with an AR-silencing lentivirus versus vector control were tested by parallel intradermal Matrigel injections into NOD/SCID male and female mice (5 per group, data of male mice in red). Mice were sacrificed 16 days after injection. A) Tumor size, measured by digital caliper (mass = (length x width x height) * π/6) together with representative low magnification H/E images of the retrieved lesions. B-E) Double immunofluorescence analysis of lesions with antibodies against AXL, for melanoma cells identification, and quantification of KI67 (B) or cytoplasmic dsDNA (C) positive cells, and mean fluorescence signal intensity of STING (D) and ICAM1 expression (E). Shown are representative images of AXL positive cells (AXL signal not shown) stained with antibodies against the other markers, together with relative quantification, (> 50 cells in 3-5 fields on digitally-retrieved images were counted using ImageJ software). F) double immunofluorescence analysis of lesions with antibodies against AXL and F4/80, for melanoma cells and macrophages identification, respectively. Shown are representative images together with quantification of the number of F4/80 positive cells per AXL positive tumor area, counting in each case 3-4 fields. Similar determination of CD45 positive cells is shown in Supplementary Figure 15. n (control versus experimental lesions) = 10; two-tailed paired t test, *P < 0.05; ** P < 0.01; ***P < 0.005. Scale bar: 10 µm. Similar tumorigenicity experiments with A375 and SKMEL28 cells plus/minus shRNA-mediated AR silencing are shown in Supplementary Figure 16, 17.

The findings are of likely translational significance, as suppression of tumor cell proliferation, together with dsDNA cytoplasmic release, STING activation and ICAM1 induction were also observed by treatment of tumor-bearing animals with the AR inhibitor AZD3514 (Figure 8A-D) or by pretreatment of cells prior to injection into the animals (Supplementary Figure 18). Even in this case, this was accompanied by increased macrophage infiltration, and engulfment of cancer cell fragments by macrophages (Figure 8E, F).

**Figure 8.**
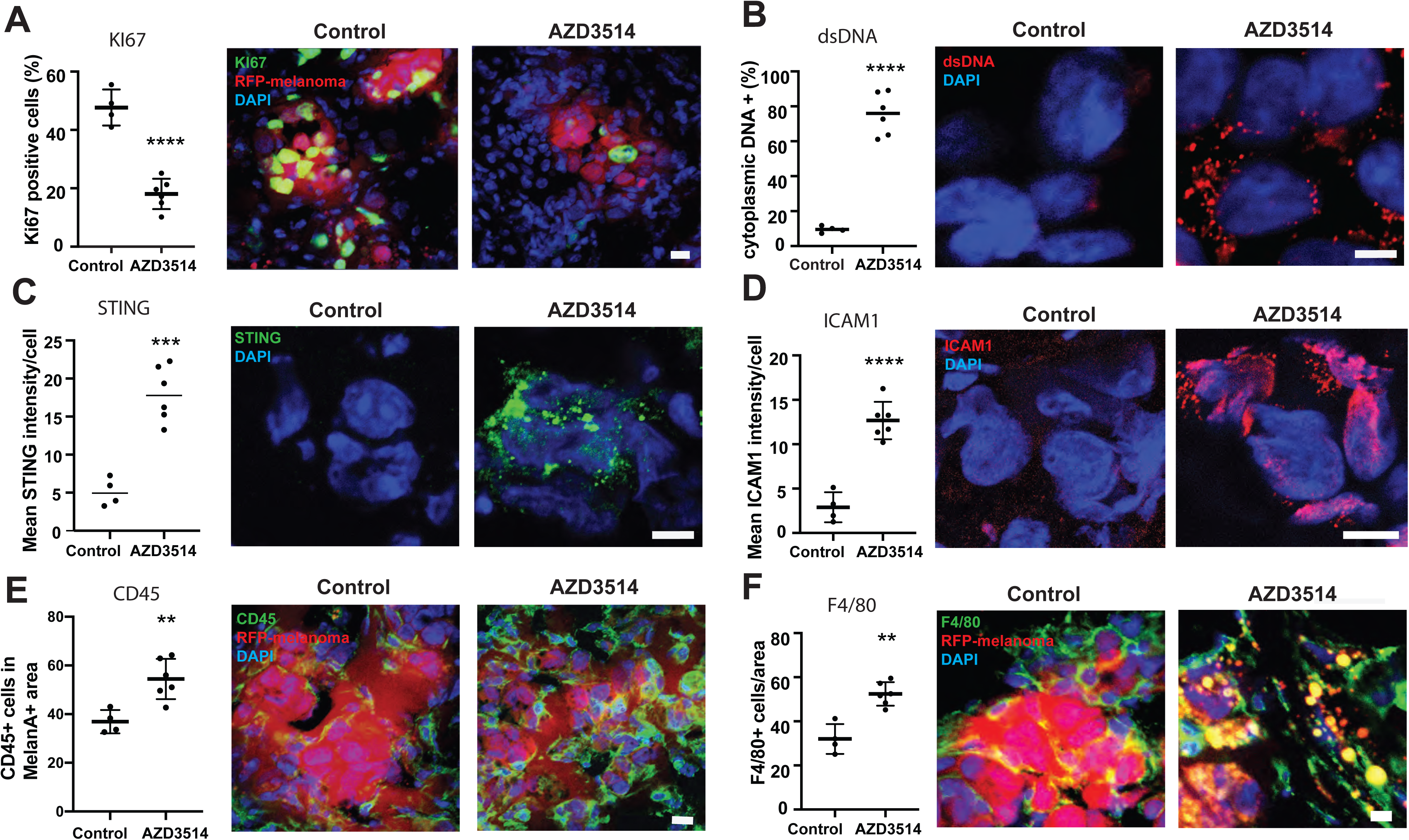
Suppression of melanoma formation by AR inhibition. RFP-expressing A375 melanoma cells were injected intradermally into 10 male mice. 3 days post-injection, mice were treated by oral gavage with either AZD3514 (50 mg/kg) or DMSO vehicle alone for 12 consecutive days. Immunofluorescence analysis was used to assess KI67 (A) and cytoplasmic dsDNA (B) positivity and STING (C) and ICAM1 (D) expression levels in melanoma cells (RFP-positive) together with numbers of juxtaposed leukocytes (E) and macrophages (F), as assessed by staining for the CD45 and F4/80 markers, respectively. Shown are quantifications together with representative images, including one (F) showing engulfment of fragmented RFP-positive melanoma cells into F4/80 positive macrophages in lesions of mice treated with the AZD3514 inhibitor. n (4 control versus 6 experimental lesions) = 10; unpaired t-test, ** P < 0.01; ***P < 0.005, ****P < 0.001. Scale bar: 10 µm (A-F). Similar tumorigenicity experiments with injection of AZD3514 pretreated WM1366 cells are shown in Supplementary Figure 18.

To further elucidate whether AR expression can influence the immunogenicity of melanoma cells and the immune infiltrates in the tumor microenvironment (40), we resorted to an immunocompetent model system based on the injection of the mouse melanoma cell line YUMM1.7 (41) into syngeneic mice (BL6 strain). Silencing of the mouse *AR* gene by two different shRNA lentiviral vectors or treatment with AR inhibitors resulted in a significant reduction of proliferation similarly to what observed with the human cells (Supplementary Figure 19).

**Figure 9.**
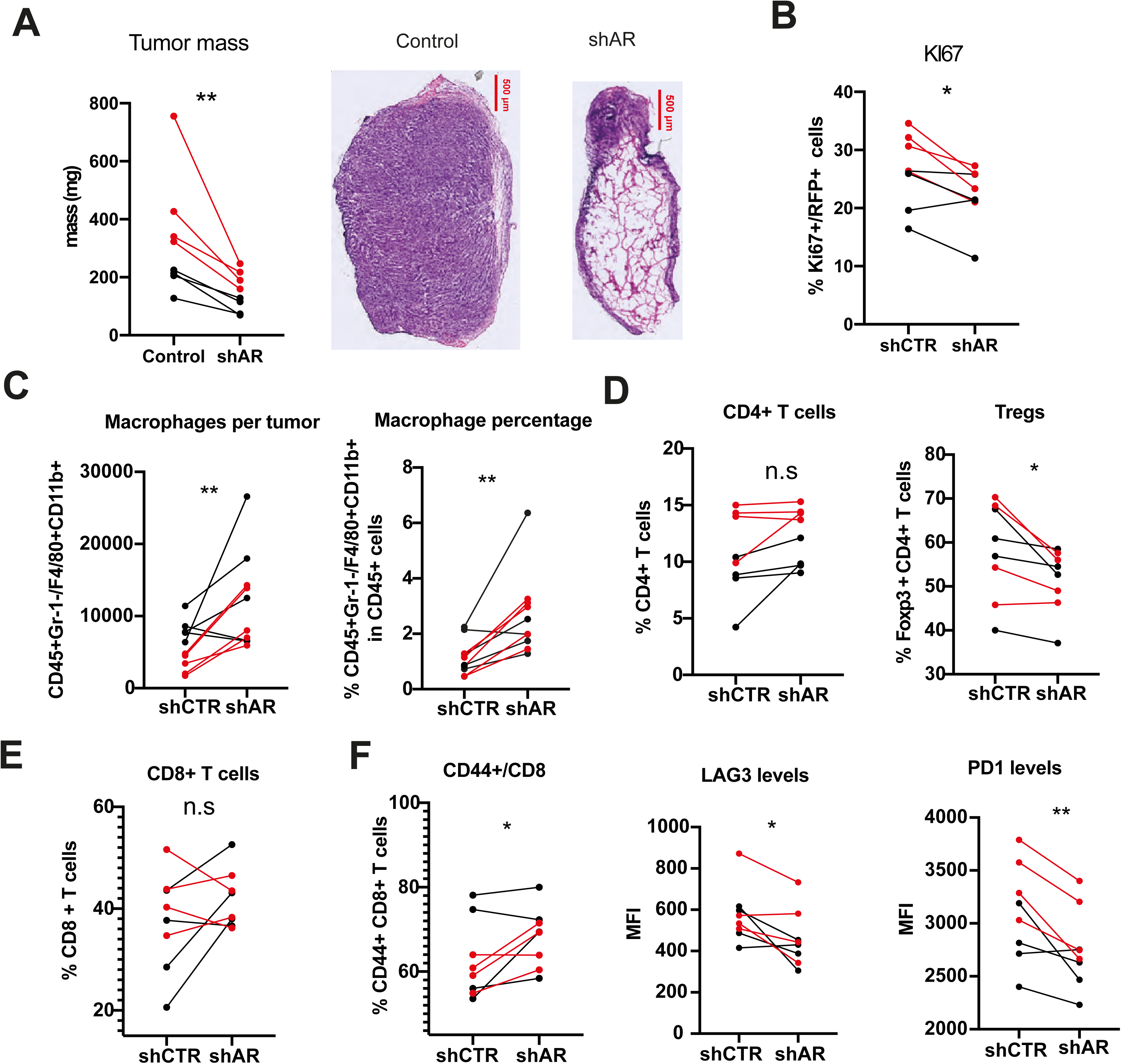
Suppression of mouse melanoma formation and immune cells recruitment by AR gene silencing. RFP-expressing YUMM1.7 mouse melanoma cells infected with an AR-silencing lentivirus versus vector control were tested by parallel intradermal Matrigel injections into BL6 male and female mice (4 per group, data of male mice in red). Mice were sacrificed 16 days after injection. A) Tumor size, measured by digital caliper (mass = (length x width x height) * π/6) together with representative low magnification H/E images of the retrieved lesions. Scale bar: 500 µm. B) Quantification of double immunofluorescence analysis for KI67 positive RPF-expressing YUMM1.7 cells in tumors. > 50 cells in 3-5 fields on digitally-retrieved images were counted using ImageJ software looking at individual cells. n (control versus experimental lesions) = 8; two-tailed paired t test, * p < 0.05. C-F) FACS analysis of tumor dissociated cells for (C) total numbers of macrophage cells (CD45^+^ Gr-1^-^ F4/80^+^ CD11b^+^) and percentage of macrophages in the CD45^+^ cell populations (left and right panels, respectively), (D) percentage of CD4^+^ T cells (CD45^+^ CD3^+^ CD4^+^) over total CD45^+^ leukocytes and fraction of TRegs (CD45^+^ CD3^+^ CD4^+^ FoxP3^+)^ within CD4^+^ T cells, (E) percentage of CD8^+^ T cells (CD45^+^ CD3^+^ CD8^+^) over total CD45^+^ leukocytes and (F) of CD44^+^ population of CD8^+^ T cells together with mean fluorescence intensity levels of LAG-3 and PD-1 staining in CD44^+^ fraction cells. n (control versus experimental lesions) = 8; two-tailed paired t test, *P < 0.05; ** P < 0.01.

*In vivo*, upon intradermal injection into immune competent mice, melanoma cells with silenced AR formed much smaller tumors than controls, with substantially reduced melanoma cell density and proliferative index (Figure 9A, B). Dissociation of tumor cells followed by FACS analysis showed a significant increase of macrophages (CD45^+^ Gr-1^-^ F4/80^+^ CD11b^+))^ in AR-silenced YUMM1.7 melanomas, consistent with what observed with human cells in immune-compromised mice (Figure 9C). While the total number of CD4^+^ T cells (CD45^+^ CD3^+^ CD4^+^) was not significantly different, that of CD4^+^ regulatory T cells (TRegs) (CD45^+^ CD3^+^ CD4^+^ FoxP3^+^) was significantly decreased in in AR-silenced YUMM1.7 melanomas (Figure 9D). Percentage levels of total CD8^+^ T cells (CD45^+^ CD3^+^ CD8^+^) did not vary consistently; however, the activated fraction (CD44^+^ population) was significantly increased with a lesser expression of co-inhibitory molecules, LAG3 and PD1, which are highly expressed by exhausted T cells (42) (Figure 9E, F). Thus, in a syngeneic mouse model, decreased tumorigenicity of melanoma cells with AR loss is associated with enhanced modulation of innate and acquired immunity.

## Discussion

The impact of sex hormone signaling in cancer development in organs with non-reproductive functions is still poorly understood (4). We have shown here that sustained AR signaling is key for melanoma cell proliferation potential and tumorigenesis in cells from male and female individuals. In addition, irrespective of its expression levels, AR plays an essential function in these cells in maintenance of genome integrity, as its genetic or pharmacologic suppression leads to genomic DNA breakage and leakage into the cytoplasm, STING activation and ensuing pro-inflammatory and immune signaling cascade. This is in contrast with previous reports on prostate cancer cells, in which AR inhibition, while synthetically lethal with other treatments (43, 44), does not appear to be sufficient to induce DNA damage and downstream events by itself. In fact, a number of reports indicate that DNA damage can be induced in prostate cancer cells by overstimulation of AR activity (45, 46).

A number of convergent mechanisms are likely to be implicated in the AR dependency of melanoma cells, involving multiple genes of the AR transcriptional signature of clinical significance that we have established. Proliferation and self-renewal potential of a large set of melanoma cells was suppressed by shRNA-mediated silencing of the gene, with similar effects resulting from down-modulation by a CRISPRi approach as well pharmacological inhibition by AR inhibitors acting through both ligand-competitive and non-competitive mechanisms. AR over-expression counteracted the shRNA gene silencing effects and was by itself sufficient to promote proliferation of melanoma cells as well as primary melanocytes with low levels of endogenous AR expression. Proliferation of these cells in charcoal-treated medium was also increased in a dose dependent manner by DHT.

We have found a 4% AR gene mutation frequency in melanoma samples of the TCGA data basis. However, the vast majority are missense gene mutations that do not coincide with those reported in the AR gene mutation data basis and their functional significance will have to be assessed. Mutations in regulatory sequences outside the AR coding region, such as those reported the TERT promoter in melanomas (47), is another interesting possibility that will have to examined. In term of predisposing genetic changes, the N terminus domain of the AR gene (NTD) contains two polymorphic trinucleotide repeats (short tandem repeat), CAG and GGN (C or T), coding respectively, for poly-glutamine and -glycine stretches of various length. The number of repeats in the AR gene has been positively or negatively associated with various cancer types, including prostate, breast and colon, even if in a number of cases the significance of this association has been an argument of contention (48). A possible association with melanoma susceptibility should be relatively straightforward to assess by combined cancer genomic and transcriptomic analysis utilizing the recently developed PCAWG platform (49).

Overall, the role that sustained AR signaling has in ensuring/promoting melanoma cells proliferation and tumorigenesis bears on the debated issue of beneficial versus detrimental consequences of androgen replacement therapy (ART) for specific aging populations, in which endogenous androgen levels diminish (50). In this context, we previously showed that in stromal fibroblasts, loss of AR function, by either gene silencing or pharmacological approaches, induces senescence of these cells together with a senescence associated secretory phenotype (SASP) that can promote tumorigenesis of neighboring cancer cells (20). Thus, inhibition of AR activity can have two-edged sword effects on cancer cells versus surrounding stromal cells, while inducing in both cellular senescence. In co-culture assays of melanoma cells and dermal fibroblasts, a potent AR inhibitor exerted net beneficial effects that paralleled those observed *in vivo* on melanoma formation in the context of the tumor microenvironment.

Besides suppression of proliferation, another major consequence of AR gene silencing or pharmacological inhibition was dsDNA breakage in the absence of additional exogenous insults, with dsDNA leakage into the cytoplasm and ensuing STING activation. The findings are consistent with the transcriptional profiles elicited by AR gene silencing in various melanoma cell lines, which are inversely associated with DNA repair gene signatures and with the pronounced down-modulation of specific genes of likely functional significance like *SENP3* gene, coding for a SUMO-specific protease with a key role in DNA repair (24, 25). While detailed mechanisms linking AR loss to DNA damage will have to be further investigated, the resulting enhancement of endogenous DNA damage suggests that already approved AR inhibitors could be used as an alternative to conventional DNA damaging agents in new combination approaches for melanoma treatment. In support of this possibility are our further findings that, in the iLINCS database, similar gene expression profiles are triggered by treatment of melanoma cells with AR inhibitors and conventional DNA damaging agents, specifically topoisomerase inhibitors.

The STING pro-inflammatory signaling cascade activated by loss of AR function can be an important determinant of tumor infiltration by immune cells (37, 51, 52). Increased cancer cells recognition and elimination by the immune system can be highly beneficial in a substantial fraction of melanoma patients (53). Bioinformatic analysis revealed that gene signatures related to interferon- and inflammatory cytokines signaling were among the most significantly associated with the gene expression profile elicited from AR gene silencing in multiple melanoma cell lines. Further EPIC (35) and CIBERSORTx (36) analysis showed that melanomas with positive association with the AR-silencing gene signature and better patients’ survival have a higher infiltration of cells of native and acquired immunity. This is consistent with our experimental findings that AR loss results in tumors with enhanced infiltration by macrophages as well as cytotoxic T cells. As such, AR inhibition could provide as an approach to ameliorate response to immune checkpoint inhibitors, especially of "immune-excluded" and "immune desert" tumors escaping innate and acquired immunity surveillance (54). This possibility will have to be carefully evaluated in the context of the complex effects that androgens as well as estrogens have on various cells of the immune system, with multiple variables including other components of the tumor microenvironment and patients’ organismic functions (55).

As suggested many years ago (9), differences in androgens levels between male and female populations are a likely determinant of their different susceptibility to the disease. However, our findings clearly indicate that melanoma cells of both male and female individuals are equally dependent on sustained AR signaling for proliferation, maintenance of genomic stability and tumorigenesis. As we have previously pointed out (4), the sexual dimorphism in this as other cancer types cannot be solely attributed to hormonal differences and/or their impact on individual cell types. In the present context, it is important to note that androgens are also produced in the female population and the AR responsive signature that we have found is equally predictive of clinical behavior in male and female melanoma patients.

At the level of individuals, the interplay between hormonal and genetic determinants of sex specification can result in a continuous spectrum of susceptibility to various diseases (56). Irrespective of sex and gender attribution, our findings point to AR signaling as a significant parameter to consider for targeted approaches to melanoma management.

## Methods

### Cell Culture

A list of different melanoma cell lines and primary melanoma cells derived from male and female patients is provided in Supplementary Table S2. Early passage (p5-6) primary melanoma cell cultures (M121008, M141022 and M131206) were established from discarded melanoma tissue samples by University Research Priority Program (URPP) Live Cell Biobank (University of Zurich) with required institutional approvals. WM1366, WM983A, WM1862, and WM1552C melanoma cells were a gift from Meenhard Herlyn (The Wistar Institute, US). The YUMM1.7 melanoma cell line was provided by Ping-Chih Ho. No further authentication of these cell lines was performed. Cell morphology and growth characteristics were monitored during the study and compared with published reports to ensure their authenticity.

All melanoma cell lines and patient-derived primary melanoma cells were maintained in Dulbecco’s modified Eagle’s medium (DMEM) (Thermo Fisher Scientific) supplemented with 10% (v/v) fetal bovine serum (Thermo Fisher Scientific) and 1% Pen-Strep.

Primary melanocytes were prepared from discarded human skin samples from abdominoplasty or circumcision at the Department of Plastic Reconstructive Surgery or Pediatrics, Lausanne University Hospital, with required institutional approvals (UNIL: CER-VD 222/12) and informed consent. All cell used in this study were determined to be negative for *Mycoplasma* prior to experiments. All cell lines were used within 5 passages after thawing.

### Cell manipulations and treatments

Lentiviral particle productions and infections were carried out as described previously (*1*). Details the lentiviral shRNA vectors and single guide RNA vectors used are provided in Supplementary Table S6. Two different shRNAs directed against human or mouse *AR* in the pLKO.1 lentiviral vector were used to silence the gene. Melanoma cells were infected with lentiviruses for 2 hours; two days post-infection cells were selected with 1 µg/ml of Puromycin for 3 days. RNA or protein samples were collected 5 days after infection. Mouse YUMM1.7 melanoma cells used for *AR* gene silencing were previously stably infected *RFP* expressing lentiviral vector with blasticidin selection.

For siRNA silencing experiments, melanoma cells were transfected with *AR* and /or *STING* silencing siRNAs versus scrambled controls siRNAs by INTERFERin® (Cat. 409, Polyplus Transfection) according to manufacturer’s instructions. The details of the siRNAs used in this study are provided in Supplementary Table 6.

For AR overexpression and rescue experiments, melanoma cells were stably infected with a blasticidin resistant lentiviral vector for constitutive *AR* expression (a gift of Karl-Henning Kalland, Bergen University, Bergen, Norway) or vector control. Post selection, the AR overexpressing melanoma cells were super-infected with an *AR* silencing or corresponding lentivirus control and selected for Puromycin resistance as described above. The cell proliferation assays were performed 5 days after the second infection.

For CRISPRi downmodulation of *AR* expression, A375 melanoma cells were stably infected with a dCas9-KRAB expressing lentivirus (pHAGE EF1a dCas9-KRAB (16)), utilizing puromycin for selection. Cells were subsequently super-infected with lentivirus (Lenti Guide-hygro-eGFP) (14) harboring scramble sgRNA or two sgRNAs targeting different regions of the *AR* promoter. Cells were analyzed 3 days after infection. The sequences of the sgRNAs are provided in the Supplementary Table 6.

For AR inhibitor treatment, 24 hours post-seeding, melanoma cells were treated with indicated concentrations of AZD3514 (Adooq Biosciences), Enzalutamide (Selleckchem) or UT155 (MedChemExpress) or DMSO solvent control as indicated. For dihydrotestosterone (DHT) treatment experiments, melanoma cells were washed 4 times in PBS after seeding and cultured for 48 hours in phenol red-free DMEM complemented with charcoal treated FBS prior to treatments with DHT (MilliporeSigma) or vehicle control (EtOH) as indicated.

### Cell based assays

*Cell proliferation assays* were carried out by measuring the production of ATP using the CellTiter-Glo luminescent assay (Promega) as per the manufacturer’s instructions. The luminescence signals for each time point were normalized to the signal obtained at day 0.

*EdU incorporation assays* were carried out using Click-iT Plus EdU Imaging Kit (Thermo Fisher Scientific) following manufacturer’s instructions. The number of EdU positive cells was analyzed and the data was represented as percentage EdU positive cells.

*Clonogenicity assays*; cells were plated on 60 mm dishes (1000 cells/well; triplicate wells/condition) and cultured for 7 days. Colonies were fixed with 4% formaldehyde and stained with 1% crystal violet. The number of clones was counted using Image J software.

*Sphere formation assays;* melanoma cells were plated onto 8-well chamber slides (Corning) pre-coated with Matrigel (Corning). In brief, chambers were coated with 100 µl Matrigel per well and incubated for 1 hour at 37 °C to polymerize. 1000 melanoma cells were plated in each well. The number of spheroids was assessed 7 days after plating through an EVOS Cell Imaging System (Thermo Fisher Scientific).

*IncuCyte cells proliferation assays*; 1000 melanoma cells per condition were seeded in triplicate into each well of a 96-well plate and allowed to attach for 12 hrs. The plates were mounted on IncuCyte Zoom System (Essen Bioscience) and cells were allowed to grow for the next 6 days. Images were captured at 4 different sectors of each well at every 2 hours for 6 days and the cell confluence was calculated by IncuCyte Zoom software.

*Apoptosis assays*; dead and pro-apoptotic cells were assessed using the Annexin Kit (BD Biosciences). In brief, before fixation, cells were washed with annexin-binding buffer, followed by incubation for 15 minutes at RT with annexin-Cy5 dye for staining of preapoptotic cells. Following annexin incubation, cell were fixed with 4% formaldehyde and counterstained with DAPI.

*Senescence-associated β-galactosidase (SA-β-Gal) activity* was assessed by the use of a commercially available chromogenic assay kit (Cell Signalling) as per the manufacturer’s instruction.

### Comet assays

The extent of double-strand DNA breaks generated with or without *AR* silencing in individual melanoma cells was assessed using alkaline comet assay (single-cell-electrophoresis) as described previously (57). Images were obtained with Zeiss AxioImager Z1. Percentage of tail DNA per nuclei was calculated using Comet Score 1.6.1.13 software (www.rexhoover.com).

### Immunofluorescence and immunohistochemistry staining

Immunofluorescences staining of tissue sections and cultured cells were carried out as described previously (35). Briefly, frozen tissue sections or cultured cells on glass coverslips were fixed in cold 4% paraformaldehyde (PFA) for 15 minutes at room temperature (RT). Paraffin embedded sections were subjected to deparaffinization and antigen retrieval using citrate-based buffer system. Samples were washed with PBS followed by permeabilization with 0.1% TritonX100 in PBS for 10 minutes and incubated with 2% bovine serum albumin in PBS for 2 hours at RT. Primary antibodies were diluted in fluorescence dilution buffer (2% bovine serum albumin in PBS, pH7.6 and incubated over night at 4°C). List of primary antibodies and dilutions used for IF is provided in Supplementary Table 6. After washing 3 times in PBS samples were incubated with donkey fluorescence conjugated secondary antibodies (Invitrogen) for 1 hour at RT. After washing with PBS, slides were mounted with Fluoromount Mounting Medium (Sigma-Aldrich) after nuclear DAPI staining. Control staining without the primary antibodies was performed in each case to subtract background and set image acquisition parameters. Immunofluorescence images were acquired with a ZEISS AxioVision or ZEISS LSM880 confocal microscope with 20X or 40X oil immersion objectives. Axiovision or ZEN Blue software were used for acquisition and processing of images. For fluorescence signal quantification, acquired images for each color channel were imported into ImageJ software, quantified using the functions “measurement” or “particle analysis” for selection of areas or cells of interest.

For the melanoma tissue microarray, the mean intensity of the AR fluorescence in melanoma cells for each micro-biopsy using ImageJ. A binary image was created from the melanA-positive cells by setting a threshold to only consider melanA signal with pixel intensity between 36-255. A mask was then derived from the melanA-positive area to mark melanoma cells. Mean intensity of AR fluorescence was then measured inside the mask. Data for melanoma tissue array data were plotted as average AR intensity of three fields per spotted tumor sample, each field comprising a group of approximately 50-60 cells. Each dot represents one clinical tissue sample.

Immunohistochemical analysis was carried out utilizing a previously described protocol for prostate cancer cells (58). Briefly, 4 μm-thick sections of formalin-fixed paraffin-embedded tissue blocks from different melanomas were subjected to de-paraffinization using xylene and hydrated in a graded series of ethanol solutions and antigen retrieval with 10 mM Tris/EDTA buffer solution (pH 9.0) at 100°C for 20 min. Parallel sections were permeabilized, blocked and incubated with anti-AR antibody or anti-melanA antibodies. Chromogenic detection was carried out using a peroxidase-conjugated secondary antibody (30 min) and DAB reagents (5 min). Tissue sections were counterstained with 0.1% hematoxylin. Immunohistochemical staining was performed by an experienced laboratory of pathology in our institution.

### Immunoblotting and qRT-PCR

Cells were lysed in RIPA buffer (10 mM Tris-Cl (pH 8.0),1 mM EDTA,1% Triton X-100, 0.1% sodium deoxycholate,0.1% SDS, 140 mM NaCl, 1 mM PMSF) or LDS buffer (Thermo Scientific). Equal amounts of proteins were subjected to immunoblot analysis. Membranes were sequentially probed with different antibodies as indicated in the figure legends, utilizing an ECL kit (Thermo Scientific) for detection. Details of antibodies used in this study are provided in Supplementary Table 6.

RT-qPCR analysis were carried out as described previously (35). A list of primers used in this study are provided in Supplementary Table 6.

### Transcriptomic Analysis

The transcriptional changes elicited in WM1366, SKMEL28 and WM115 melanoma cells plus/minus *AR*-silencing with two different lentiviruses versus empty vector control were assessed by Clariom^TM^ D GeneChip array analysis (Thermo Fisher Scientific). 5 days post-infection, RNA was extracted from the melanoma cells using Direct-zol RNA MiniPrep kit (Zymo Research) coupled with DNase treatment and RNA quality was verified by Bioanalyzer (Agilent Technologies). 50 ng of total RNA was used as input for the preparation of single-strand cDNA using the GeneChip WT PLUS Reagent Kit (Thermo Fisher Scientific). Targets were then fragmented and labeled with the GeneChip WT Terminal Labeling Kit (Thermo Fisher Scientific) and hybridized on Human ClariomTM D GeneChip arrays (Thermo Fisher Scientific) at the iGE3 Genomics Platform, University of Geneva (Geneva, Switzerland). Data obtained were analyzed using the TAC software (v4.0). The data generated in this study has been deposited to the public functional genomics data repository GEO (Gene Expression Omnibus), NCBI with an accession number GSE138486.

Gene set enrichment analysis (GSEA) for GeneChip microarray data were conducted using GSEA software using default parameters. Curated gene sets were obtained from the Molecular Signatures Database (MSigDB version 5.2, www.broadinstitute.org/gsea/msigdb/). A list of enriched pathway gene sets is provided in Supplementary Table S4.

### Construction of the *AR* silencing gene signature

Raw microarray expression data were preprocessed with TAC software, obtaining gene-level expression values from SST-RMA summarization. Ensemble IDs were mapped to gene symbols with Biomart. A paired differential expression analysis between control (n=3) and shAR (n=6) conditions was performed with Limma (default parameters), pairing together samples from each cell line. The AR silencing signature was constructed as the list of genes up- or down-regulated upon AR silencing, i.e. genes showing an adjusted P-value < 0.05 and an absolute log-fold change > 1 in the overall analysis as well as in each cell line separately.

### Computation of AR silencing signature scores in TCGA SKCM dataset

Level 3 gene expression and clinical data for Skin Cutaneous Melanoma (SKCM) TCGA projects were downloaded from NIH GDC Data Portal (https://portal.gdc.cancer.gov). In case both primary and metastatic samples were present for the same patient, only the primary sample was retained. TPM (transcripts per million) values were transformed as log2 (TPM+1). Scores for the sets of genes up- and down-regulated were computed with the R package ‘GSVA’ v1.30.0 using default parameters. The difference between these scores was computed to obtain a unified score for the total *AR* silencing gene signature (comprising both up- and down-regulated genes) for each patient. Each tumor was assigned with a positive (+) or negative (-) score relative to the unified *AR* silencing gene signature.

### Survival analysis

The difference between the survival of patients with a positive (+) versus negative (-) score for the *AR* silencing gene signature was tested with a log-rank test implemented in the R package ‘survival’ v2.43-3 (https://CRAN.R-project.org/package=survival). A Cox regression from the same package was used to account for the following covariates: age, sex, primary or metastatic status and genomic subtype (BRAF mutant, RAS mutant, NF1 mutant or triple wild-type).

### EPIC and CIBERSORTx analyses

Cell type fractions for bulk RNA-seq melanoma samples from TCGA SKCM were computed with EPIC v1.1.5 (http://epic.gfellerlab.org) using default parameters and CIBERSORTx (https://cibersortx.stanford.edu/) using LM22 signature matrix and “B-mode” batch correction. Differences between melanomas enriched and not enriched for the *AR* silencing gene signature were tested with Wilcoxon rank-sum tests.

### iLNCS analysis

The analysis of concordance between our in-house *AR* silencing gene signature and iLINCS chemical perturbagen signatures was performed by interrogating the iLINCS data portal (http://www.ilincs.org/ilincs/). Briefly, the iLINCS web application computes concordance as the Pearson correlation coefficient between the fold changes of the genes in common (n=21) between the query signature and the pre-computed iLINCS signatures. Signatures with correlation > |0.2| and p-value < 0.05 are extracted and sorted by concordance. A list of top iLINCS signatures with concordance score >0.65 is shown in Supplementary Table S5.

### Tumorigenesis experiments

Intradermal back injections of indicated melanoma cells were carried out in 6 to 8-week-old male and female NOD SCID mice (NOD.CB17-Prkdcscid/J, Jackson Laboratory). In brief, 1 x 10^6^ melanoma cells (WM1366, A375, SKMEL28) infected with *AR* silencing vs control lentiviruses were injected with Matrigel (Corning) (70 µl per injection) intradermally in parallel into the left and right side of mice with 29-gauge syringes. Mice were sacrificed and Matrigel nodules were retrieved for tissue analysis 16 days after injection.

For *in vivo* AZD3514 treatment experiments, RFP-expressing A375 melanoma cells (1 × 106) were intradermally injected with Matrigel solution in the back skin of 10 male NOD SCID mice. Three days post-injection mice were either treated with 100 µl of AZD3514 (50 mg/kg, per mouse, group of 5 mice) or with Captisol (Ligand Technology) as vehicle control (a group of 4 mice) for consecutive 12 days by oral gavage. The bodyweight of the mice was measured regularly during the treatment. Mice were sacrificed and Matrigel nodules were retrieved for tissue analysis at the end of the treatment.

Alternatively, melanoma cells were either treated with AZD3514 (10 µM) or DMSO for 12 hrs in culture and injected intradermally in mice together with Matrigel as described above. The tumors were allowed to grow for two weeks and nodules were retrieved for tissue analysis at the end of the treatment.

Intradermal back injections of YUMM1.7-RFP cells were carried out in 6 to 8-week-old male and female mice (C57BL/6J, Jackson Laboratory). In brief, 5 x 10^5^ melanoma cells infected with *AR* silencing vs control lentiviruses were injected with Matrigel (Corning) (60 µl per injection) intradermally in parallel into the left and right side of mice with 29-gauge syringes.

Mice were sacrificed and Matrigel nodules were retrieved for flow cytometry analysis 14 days after injection. All mice were housed in the animal facility of the University of Lausanne.

### Tumor digestion, cell isolation and flow cytometric analysis

Tumors were minced in RPMI with 2% FBS, intravenous collagenase (0.5 mg/ml, Sigma-Aldrich) and DNase (1 µg/ml, Sigma-Aldrich) and digested at 37°C for 45 min. The digested samples were then filtered through a 70 µm cell strainer and washed with fluorescent activated cell sorter buffer (phosphate buffered saline with 2% fetal bovine serum and 2 mM EDTA). The cell pellets were then incubated with ACK lysis buffer (Invitrogen) to lyse red blood cells. Next, viable cells in single-cell tumor suspensions were further enriched by density gradient centrifugation (800g, 30 min) at room temperature with 48% and 80% percoll (GE healthcare) and collected from the interphase of the gradient. Fluorescent activated cell sorter analysis was performed using LSRII (BD Biosciences). Data were analyzed using FlowJo. The following antibodies were used for flow cytometry: anti-CD3 (17A2), anti-CD4 (RM4-5), anti-CD8*α*(53-6.7), anti-CD11b (M1/70), anti-CD45 (104), anti-Gr-1 (RB6-8C5), anti-FoxP3 (FJK-16s), anti-CD44 (IM7), anti-F4/80 (BM8), anti-LAG3 (C9B7W), anti-PD1 (RMP1-30) and anti-Arg1 (A1exF5). Cell populations were identified based on the expression markers listed here.

CD4 T cells: CD45^+^/CD3^+^/CD4^+^; CD8 T cells: CD45^+^/CD3^+^/CD8^+^; T_reg_s: CD45^+^/CD3^+^/CD4^+^/FoxP3^+^; Macrophage: CD45^+^/Gr-1^-^/F4/80^+^/CD11b^+.^ Antibodies details - including commercial sources - are provided in Supplementary Table 6.

### Statistical analysis

Statistical testing was performed using Prism 7 (GraphPad Software). Data are presented as mean± SEM or mean ± SD, as indicated in the legends. Statistical significance for comparing two experimental conditions was calculated by two-tailed t-tests. For multiple comparisons of more than two conditions, one-way ANOVA was employed, with Dunnett’s test to compare different test conditions to the same control. For tumorigenicity assays, wherever possible, individual animal variability issue was minimized by contralateral injections in the same animals of control versus experimental combinations of cells. No statistical method was used to predetermine sample size in animal experiments and no exclusion criteria were adopted for studies and sample collection.

### Study approvals

Melanocytes were prepared from discarded human skin samples from abdominoplasty or circumcision at the Department of Plastic Reconstructive Surgery or Pediatrics, Lausanne University, with required institutional approvals (UNIL: CER-VD 222/12) and informed consent. Benign nevi, dysplastic nevi, primary and metastatic skin sections, and melanoma tissue microarray slides were obtained from the Live Cell Biobanks of the University Research Priority Program (URPP) “Translational Cancer Research” (Mitchell P. Levesque, University Hospital Zurich). All samples were obtained as surplus material from consenting patients (Ek.647/800), and the experiments were approved by the Kantonal ethical committee of Zürich (Kantonale Ethikkommission Zürich, Zürich, Switzerland, approval no. KEK.Zh.Nr.2014-0425). No access to sensitive information has been provided.

All animal studies were carried out according to Swiss guidelines for the use of laboratory animals, with protocols approved by the University of Lausanne animal care and use committee and the veterinary office of Canton Vaud (animal license No. 1854.4f/ 1854.5a).

## Author contributions

MM, SG, AC, GB, AK, LM, JE, AS performed the experiments and/or contributed to analysis of the results. DT and GC performed the bioinformatics analysis. YRY and PCH performed the FACS analysis. ML and RD provided clinical samples. BO helped with the writing of the manuscript. MM, SG and GPD designed the study, assessed the data and wrote the manuscript. MM and SG share co-first authorship based on their contribution to the work.

## Acknowledgments

We thank Meenhard Herlyn for the gift of WM1366, WM983A, WM1862, WM1552C cells; Pascal Schneider for the gift of STING antibody; Fabio Martinon for stimulating discussions; Xiaoyun Li, Haiping Wang, Beatrice Tassone and Tatiana Proust for technical help. This study was supported by grants from the Swiss National Science Foundation (310030B_176404 “Genomic instability and evolution in cancer stromal cells”), the Swiss Cancer League (KFS 4709-02-2019), the European Research Council (26075083), and the NIH (R01AR039190, R01AR064786; the content does not necessarily represent the official views of the NIH). We thank the University Research Priority Program (URPP) in Translational Cancer Research Biobank at the University of Zürich for access to the melanoma cell lines.

## Supplementary Materials

**Supplementary Figure 1.**
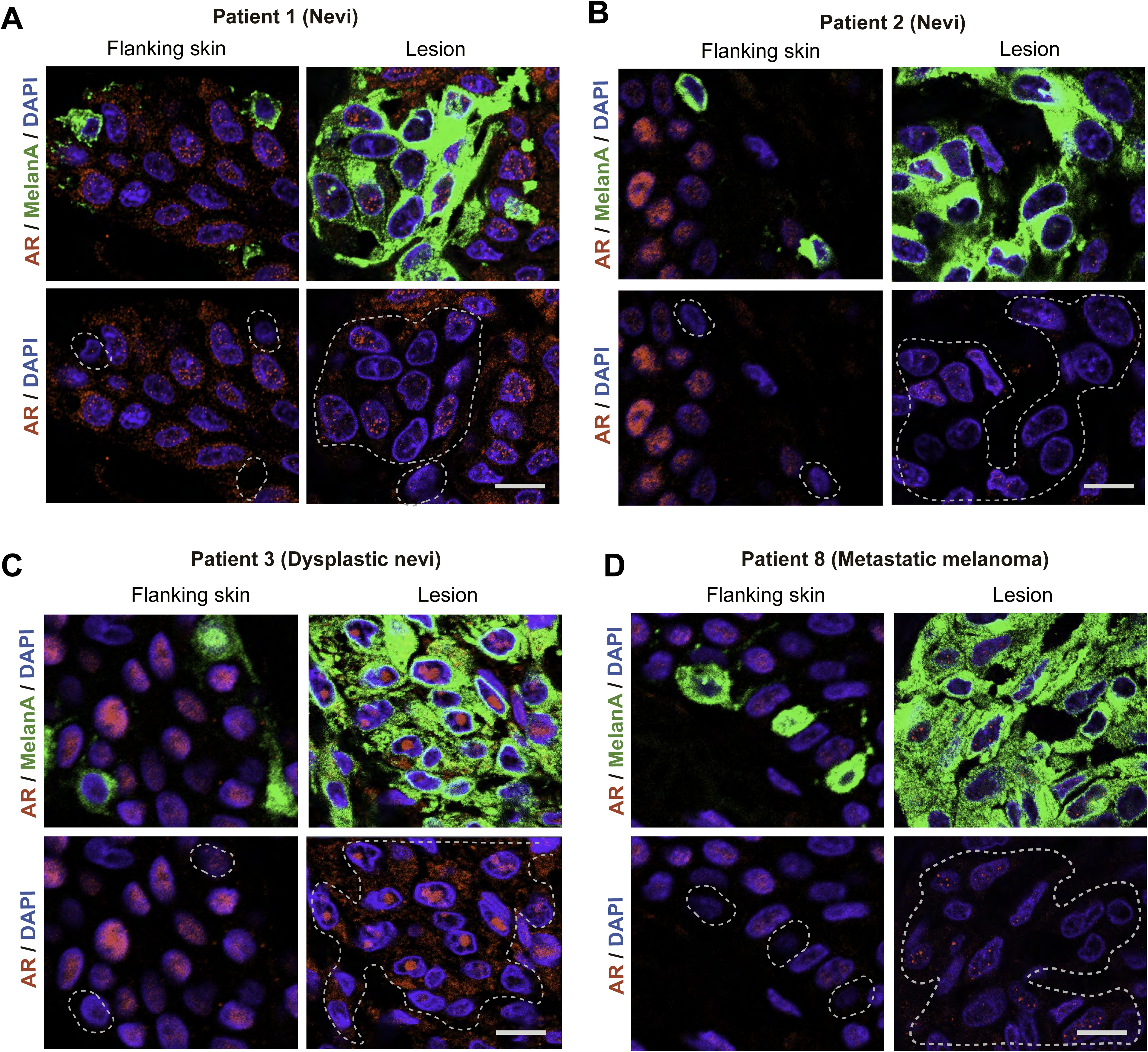
Related to Figure 1A. Double immunofluorescence analysis of patient-derived melanocytic lesions. Double immunofluorescence images of benign nevi (A, B), dysplastic nevi (C), and metastatic melanoma (D) in parallel with flanking skin stained with anti-MelanA (green) and anti-AR (ab74272) (red) antibodies. Highlighted in the lower panels are representative MelanA positive cells and areas used for quantification in Figure 1A. Scale bar: 10 µm.

**Supplementary Figure 2.**
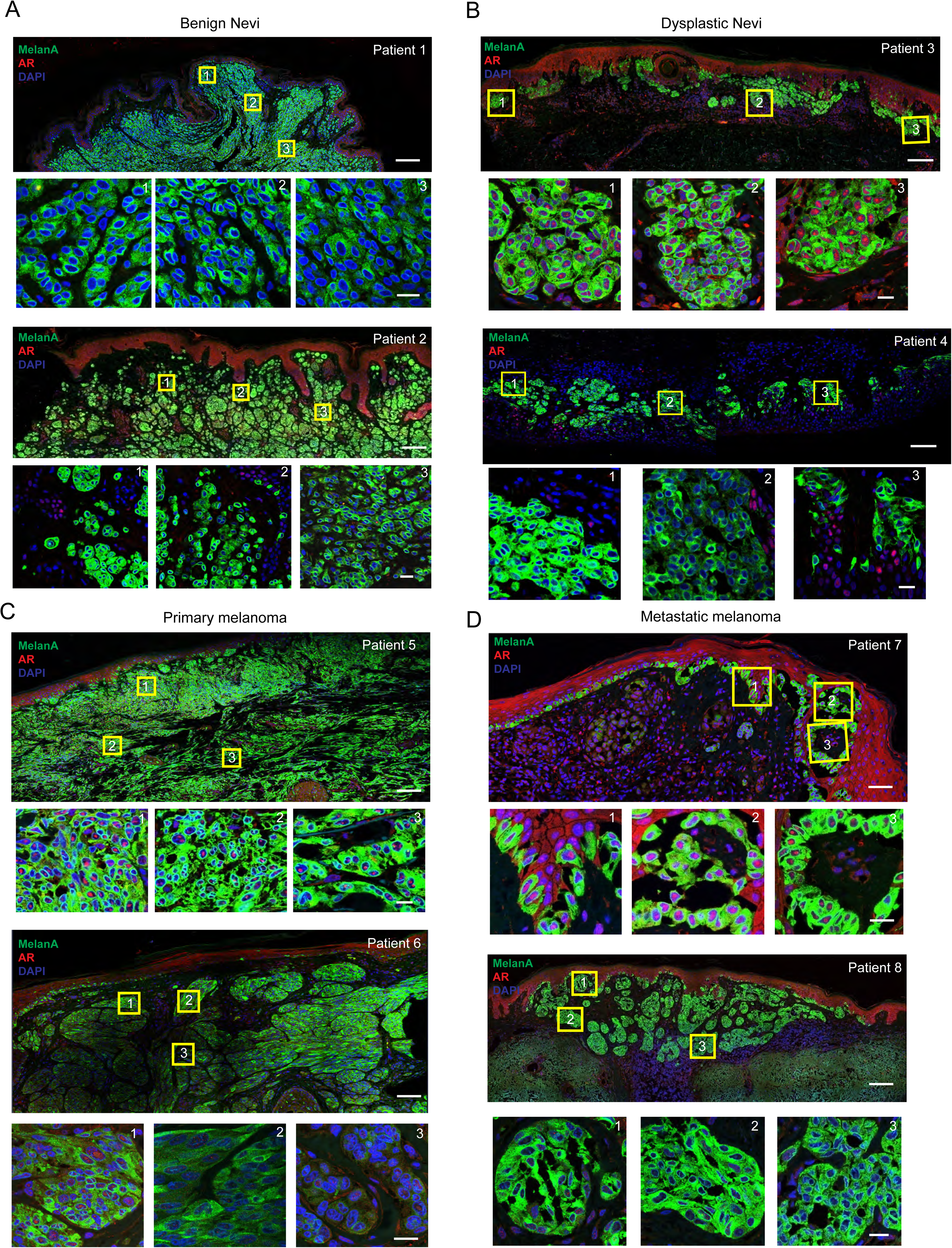
Related to Figure 1B. Double immunofluorescence analysis of patient-derived melanocytic lesions. Immunofluorescence staining of benign nevi (A, patient 1 and 2), dysplastic nevi (B, patient 3 and 4), primary melanoma (C, patient 5 and 6) and metastatic melanoma (D, patient 7 and 8) skin tissues with anti-MelanA (green) and anti-AR(ab74272) (red) antibodies, and topographically distinct areas (boxes 1, 2 and 3) utilized for single cell AR expression quantification in Fig. 1B. Shown are representative low and high magnification images of the areas used for quantification. Scale bar: 2 mm and 20 µm, respectively.

**Supplementary Figure 3.**
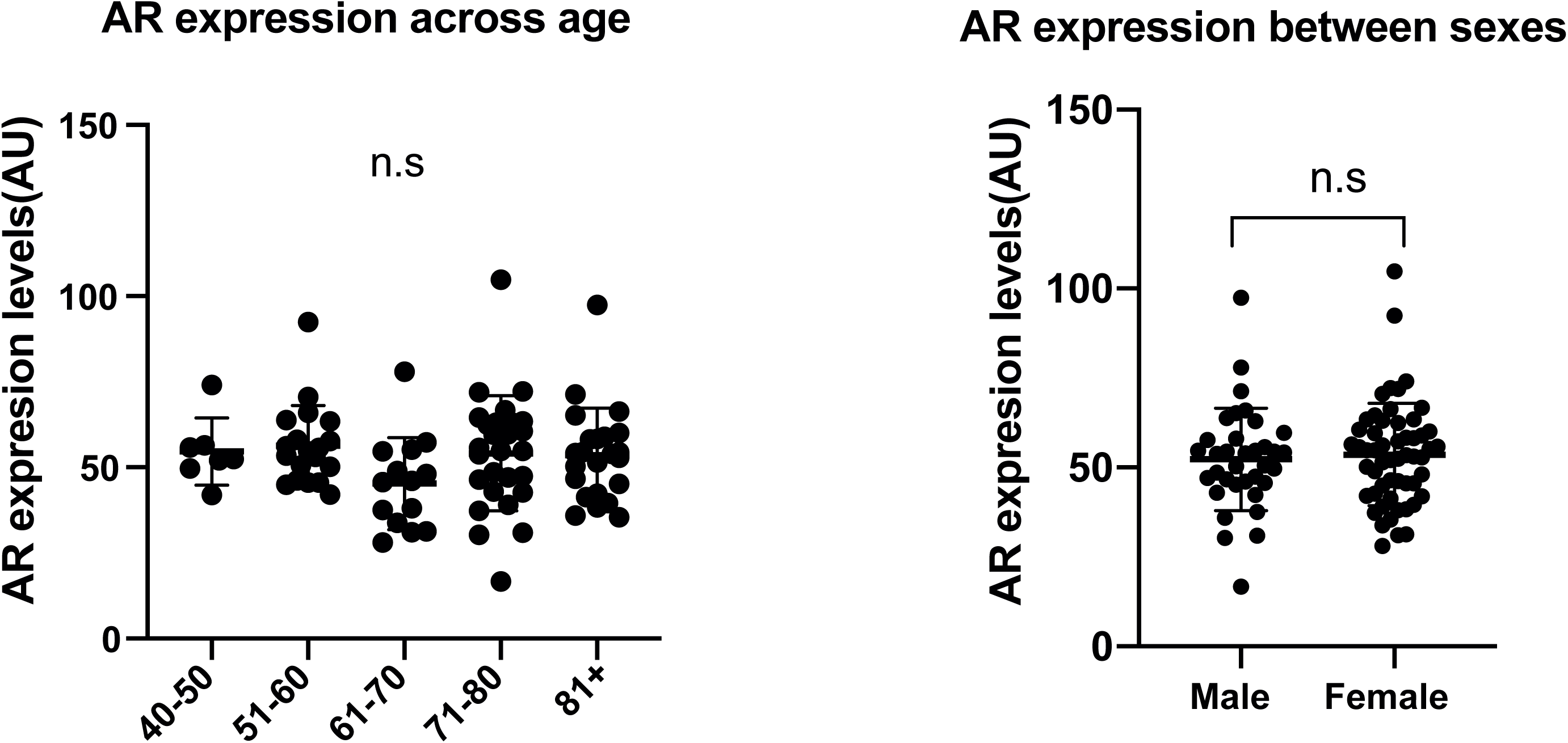
Related to Figure 1C. AR expression across age and between sexes in a melanoma tissue microarray. Quantification of AR fluorescence signal in MelanA-positive cells in a tissue microarray of melanoma patients divided by age or sex. Quantification was based on digitally-acquired images of three independent fields per clinical lesion (a minimum of 50 cells per field) on the arrays. Results are expressed as average values for each lesion (dots) together with mean across years of age (left) or sex (right) of patients.

**Supplementary Figure 4.**
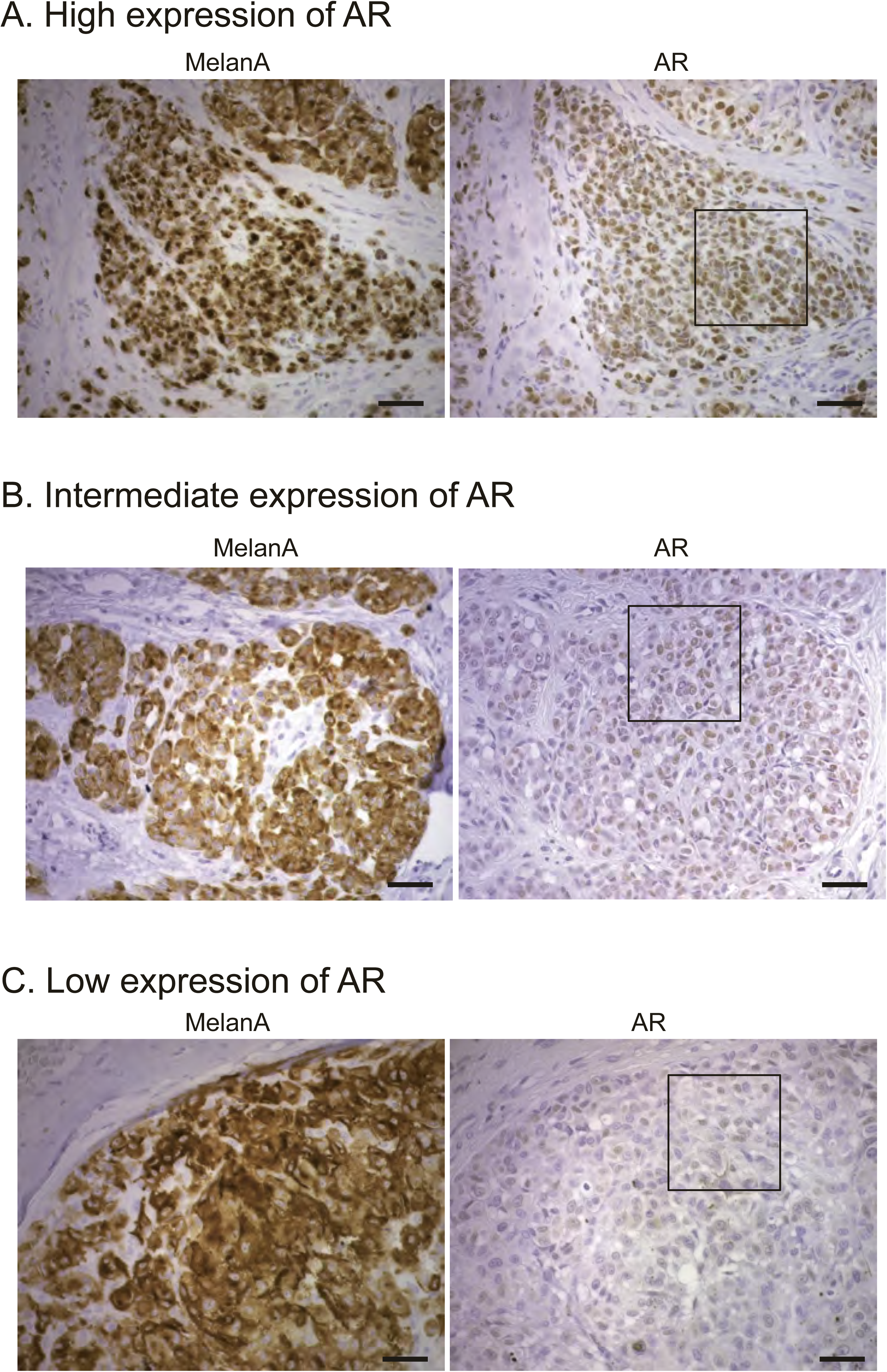
Related to Figure 1D. Immunohistochemical analysis of AR expression in patient-derived melanocytic lesions. Immunohistochemical staining with anti-MelanA and anti-AR (ab74272) antibodies of parallel sections of different melanomas with high (A), intermediate (B) and low (C) level of AR expression as quantified by double immunofluorescence analysis in Fig. 1B. Shown are representative images, with enlarged boxed areas shown in Fig. 1D. Scale bar: 50 µm

**Supplementary Figure 5.**
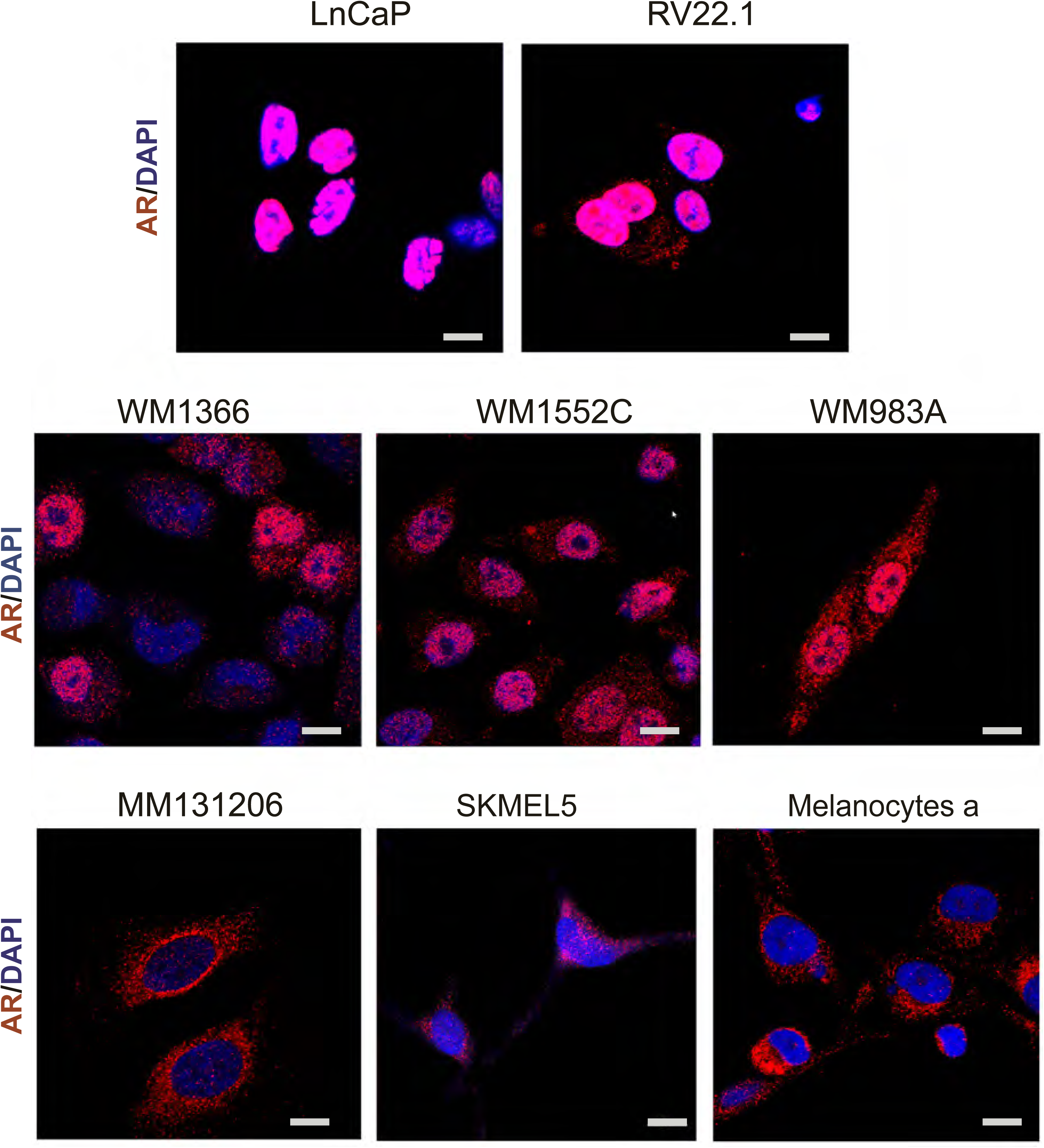
Related to Figure 1E. Immunofluorescence analysis of AR expression in different melanoma cell lines and primary human melanocytes with prostate cancer cells as comparison. Representative images of the indicated prostate cancer cells lines (LnCaP, 22RV.1), melanoma cell lines and primary melanoma cells with high (WM1366, WM1552C, WM983A) and low AR (MM131206, SKMEL5) expression and primary human melanocytes (strain a) stained with anti-AR (red) antibody (D6F11) and DAPI (blue) nuclear staining. Scale bar: 10 µm.

**Supplementary Figure 6.**
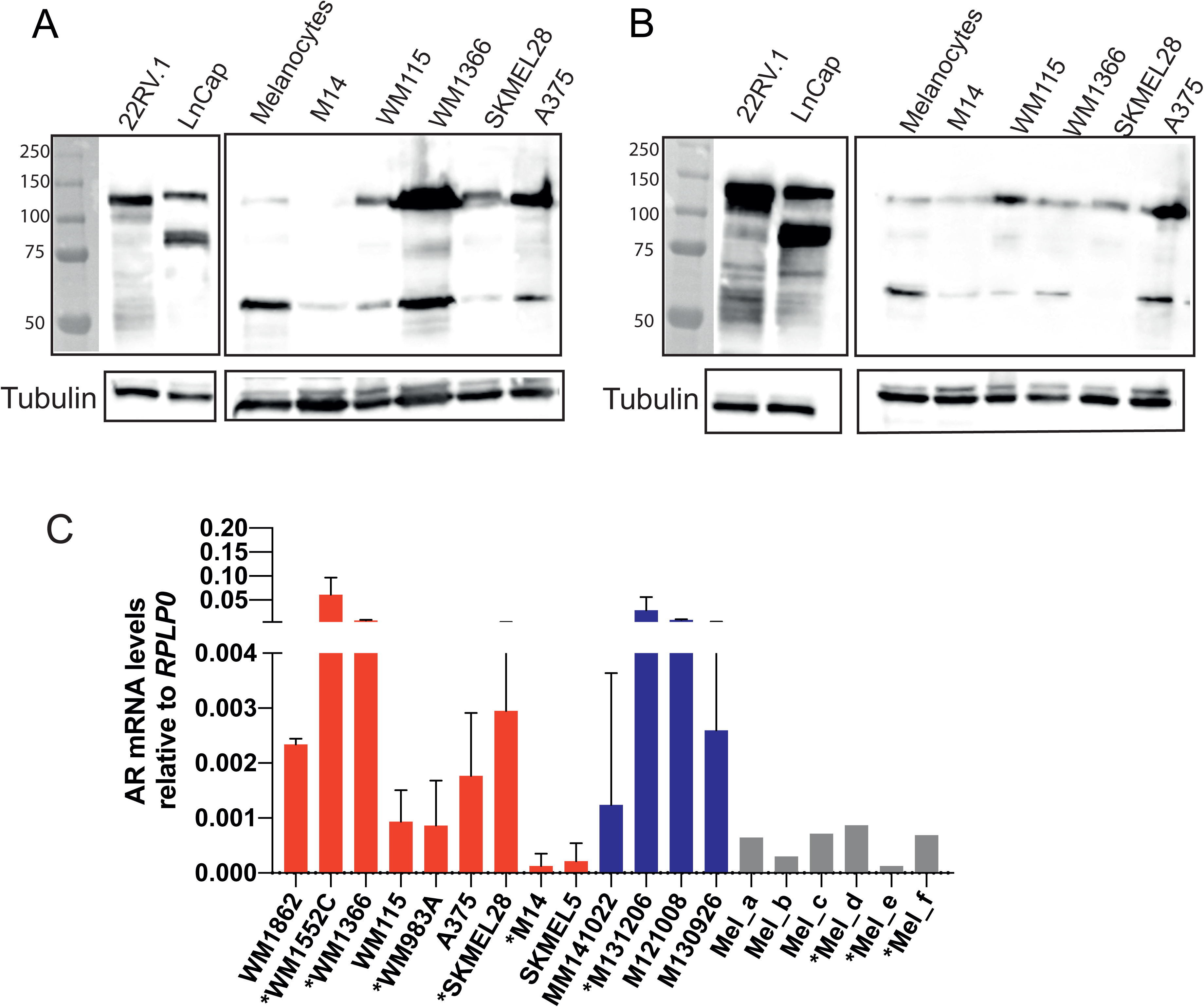
Related to Figure 1. AR expression in different melanoma cell lines and primary human melanocytes as detected by two different antibodies. AR expression) in melanoma cell lines (A375, SKMEL28, WM1366, WM115 and M14) and primary human melanocytes was assessed by immunoblot analysis with two different antibodies in parallel with prostate cancer cell lines (LnCaP, 22RV.1) as comparison. All extracts were run in two parallel gels and blotted, respectively, with anti-AR (D6F11) (A) or anti-AR (PG-21) antibodies (B). Shown are low and high exposure images of the same blots, for better AR detection in highly expressing prostate cancer versus melanoma cells. Note a band of the expected molecular size for full length AR proteins (110 KD) detected in all tested cells and a second band around 60KD detected by the two antibodies, as a possible product of proteolytic cleavage. C) RT-qPCR analysis of AR mRNA expression in a panel of melanoma cell lines (red), early passage primary melanoma cells (blue) and primary human melanocytes (grey). Results are expressed as relative to RRLP0 values.

**Supplementary Figure 7.**
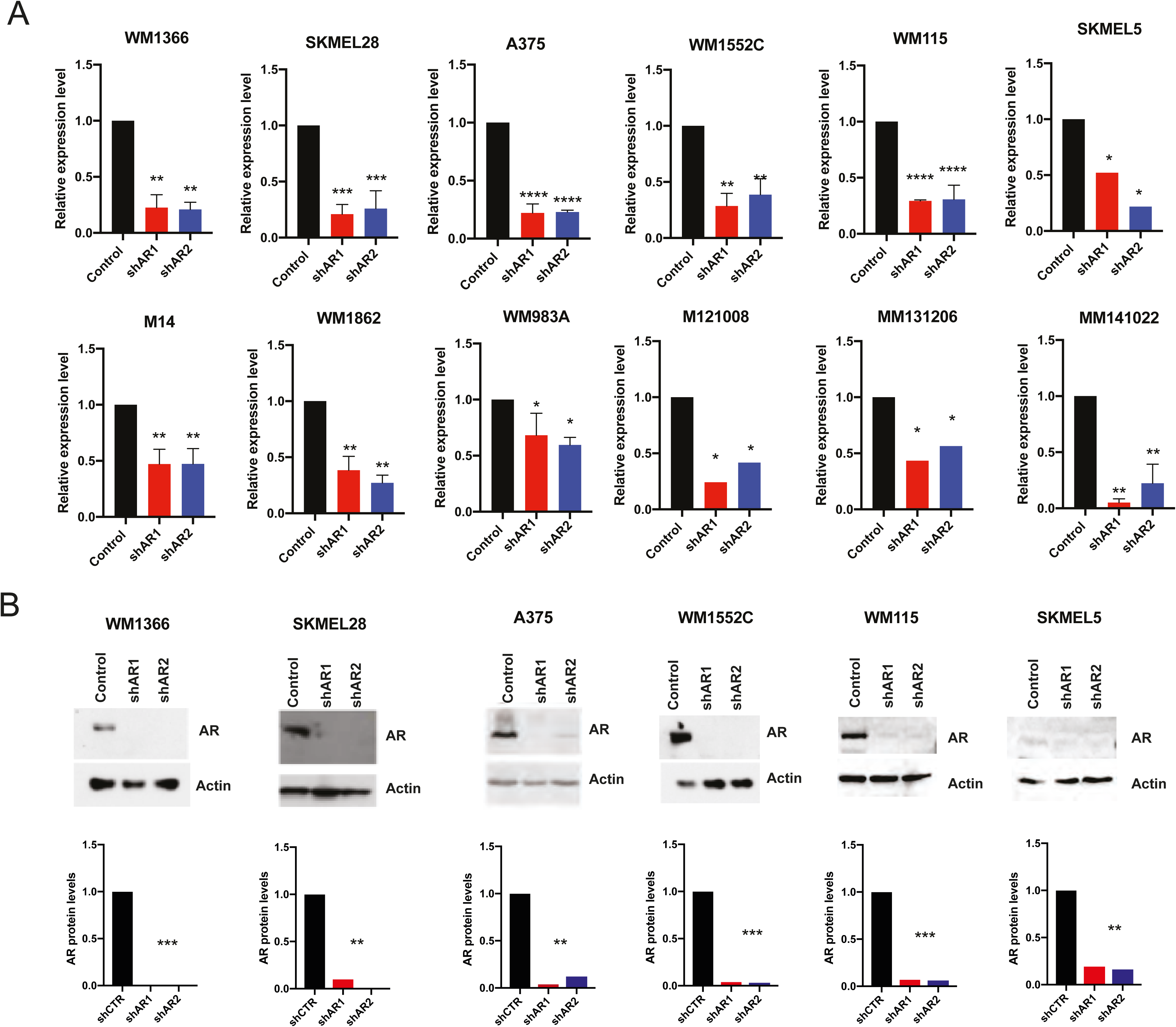
Related to Figure 2A-F. Silencing of AR in different melanoma cell lines. A) Down-modulation of AR expression in a panel of melanoma cell lines and primary melanoma cells (M121008, MM131206, MM141022) infected with 2 AR silencing lentiviruses versus empty control (5 days after infection) was assessed by RT-qPCR. Data are shown as mean ± SD, 1-way ANOVA with Dunnett’s test. n = 3 biological replicates (experiments). *P < 0.05; ** P < 0.01; ***P < 0.005; ****P < 0.001. B) Immunoblot analysis of AR protein expression in different melanoma cell lines plus/minus AR gene silencing as in previous panel. Shown are the immunoblots together with the corresponding quantification of AR protein levels after densitometric scanning of the autoradiographs, utilizing actin signal for normalization (lower panels).

**Supplementary Figure 8.**
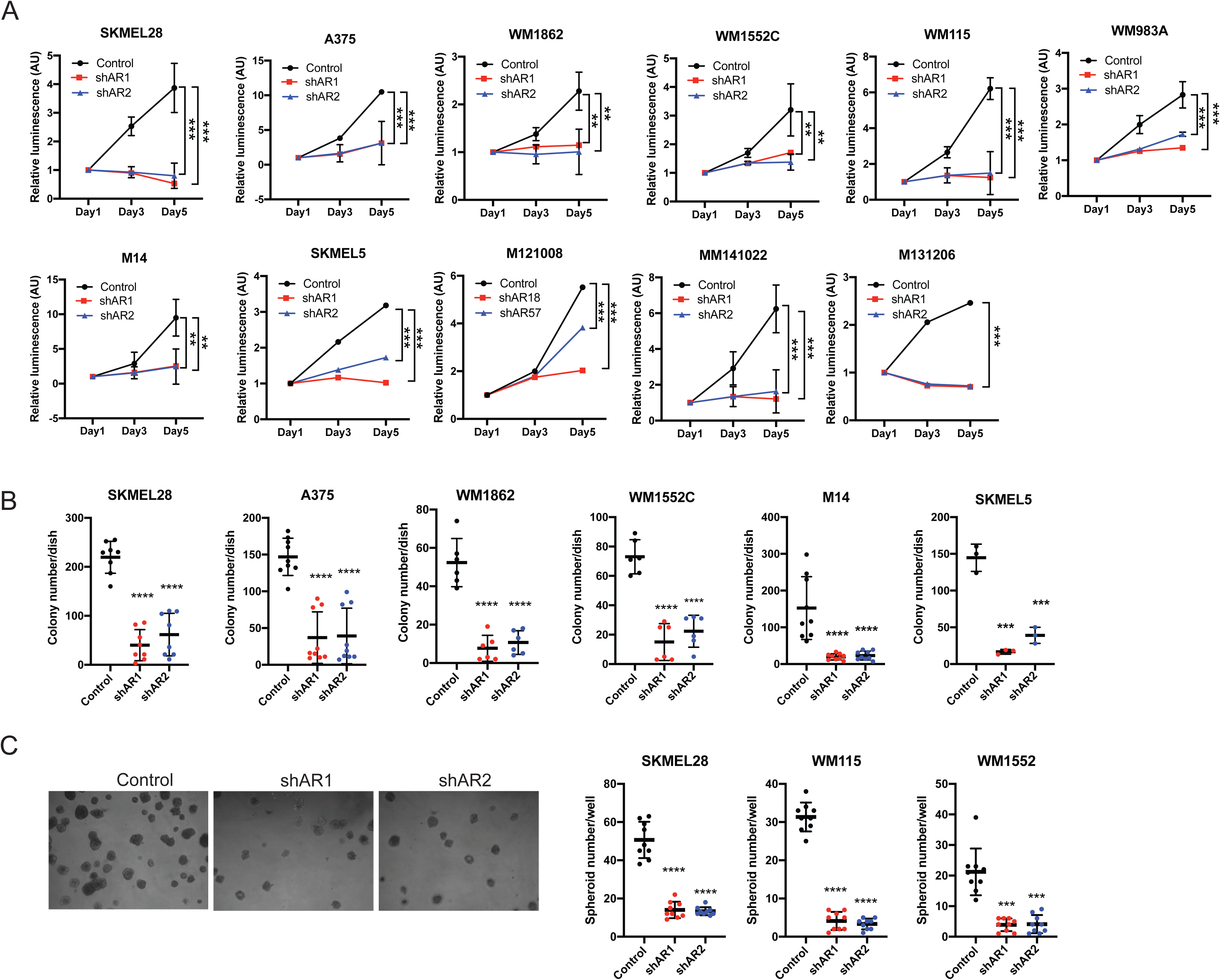
Related to Figure 2A-C. Suppression of melanoma proliferation and self-renewal potential by AR silencing. A) Cell density assays (CellTiter-Glo) were carried out with the indicated melanoma cell lines and primary melanoma cells (M121008, MM131206, MM141022) infected with two AR silencing lentiviruses versus empty vector control. Results are presented as luminescence intensity values relative to day 1. B, C) Colony and sphere formation assays with indicated melanoma cell lines plus/minus AR silencing. Shown are the results of 3 independent experiments quantifying in each case 3 dishes per conditions (indicated by dots, mean ± SD). Results are presented as mean ± SD, 1-way ANOVA with Dunnett’s test. n = 3 biological replicates (experiments). ** P < 0.01; ***P < 0.005; **** <0.001.

**Supplementary Figure 9.**
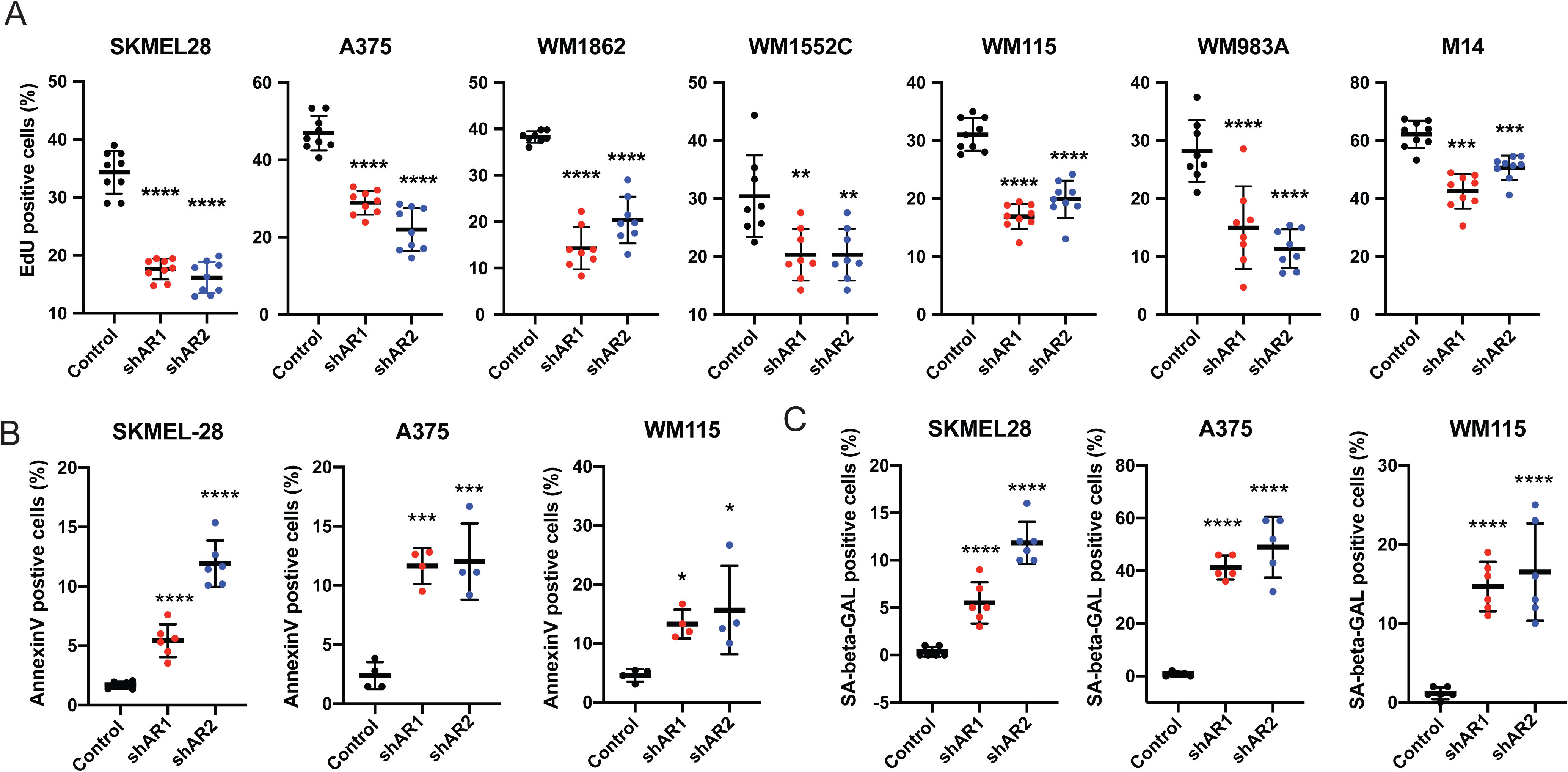
Related to Figure 2D-F. EdU incorporation, apoptosis and senescence assays in melanoma cells plus/minus AR silencing. Indicated melanoma cell lines infected with two AR silencing lentiviruses versus empty vector control were tested by EdU labelling assay (A), AnnexinV staining (B) and senescence-associated beta-GAL staining (C) 5 days post virus infection. AnnexinV/SA-beta-GAL -positive cells were counted using ImageJ software. Shown are representative images and results of 3 independent experiments quantifying in each case 3 dishes per conditions (indicated by dots, mean ± SD), 1-way ANOVA with Dunnett’s test. n = 3 biological replicates (experiments). *P < 0.05; ** P < 0.01; ***P < 0.005, ****P < 0.001.

**Supplementary Figure 10.**
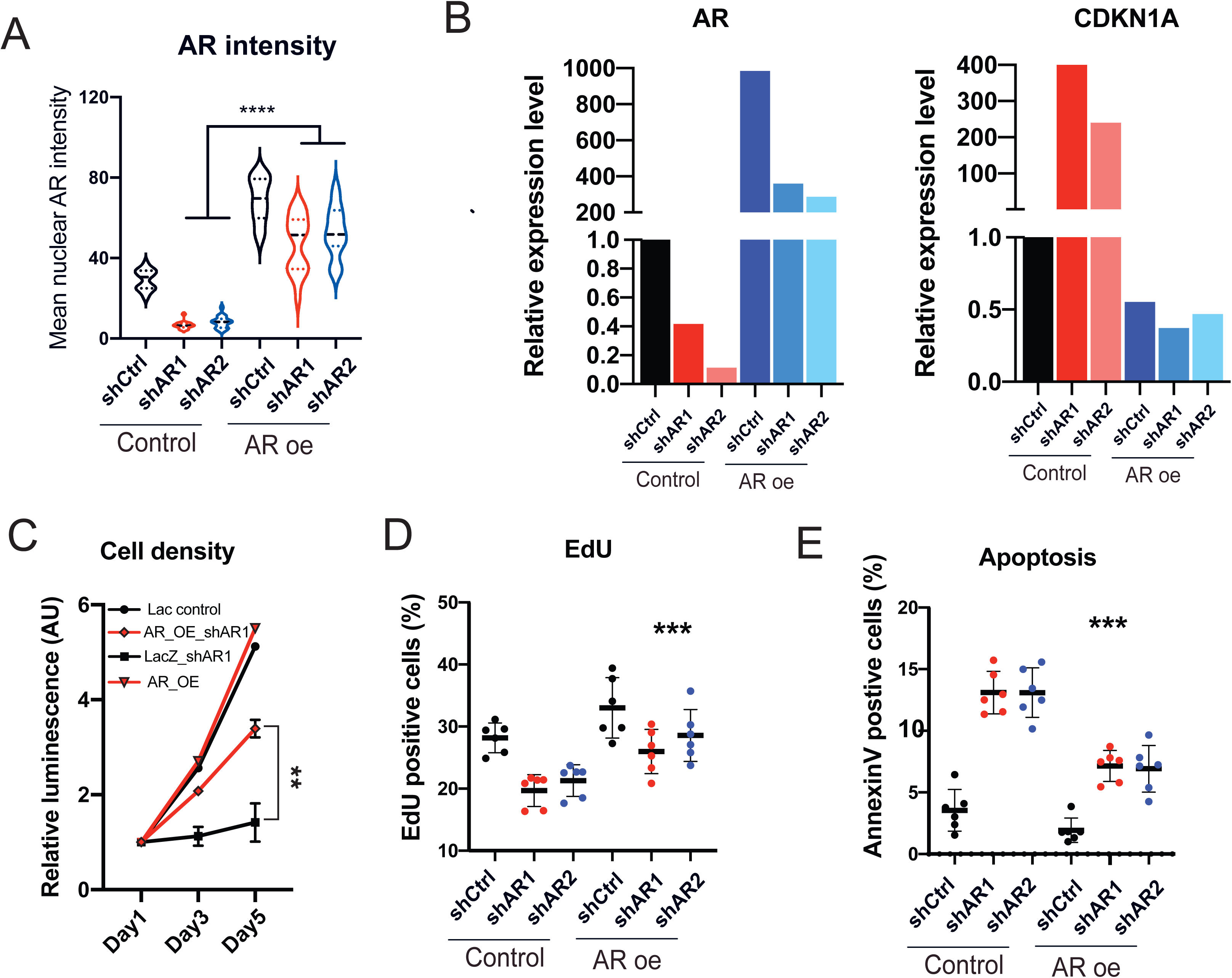
Related to Figure 2G. Concomitant AR overexpression suppresses AR silencing effects. A) Quantification of AR protein expression by immunofluorescence analysis of A375 cells stably infected with a lentiviral vector for constitutive AR expression versus LacZ control and superinfected with two AR silencing lentiviruses versus vector control for 5 days. Shown is a violin plot quantification of AR immunofluorescence signal intensity, with corresponding representative images shown in Fig. 2G. n (cells per condition) > 20, 1-way ANOVA with Dunnett’s test, ****P < 0.001. B) Quantification of AR mRNA expression by RT-qPCR analysis of A375 cells plus/minus AR overexpression and silencing as in the previous panel. Same samples were analyzed for levels of *CDKN1A* expression as a marker/effector of cellular senescence induced by AR gene silencing. C) Proliferation live-cell imaging assays of A375 cells plus/minus AR overexpression and silencing as in the previous panel. Cells were plated in triplicate wells in 96-well plate followed by cell density measurements ((IncuCyteTM system, Essen Instruments; 4 images per well every 4 hours for 128 hours). n (number of wells) = 3, Pearson r correlation test. ** P < 0.01. D, E) Same melanoma cell as in the previous panels were tested by EdU labelling assay (D) or apoptosis by annexin V staining (E). For each condition, cells were tested in duplicated dishes, with all experiments repeated 3 times. Data are shown as mean ± SD, 1-way ANOVA with Dunnett’s test. n = 3 independent experiments. **P < 0.01. *** P<0.005.

**Supplementary Figure 11.**
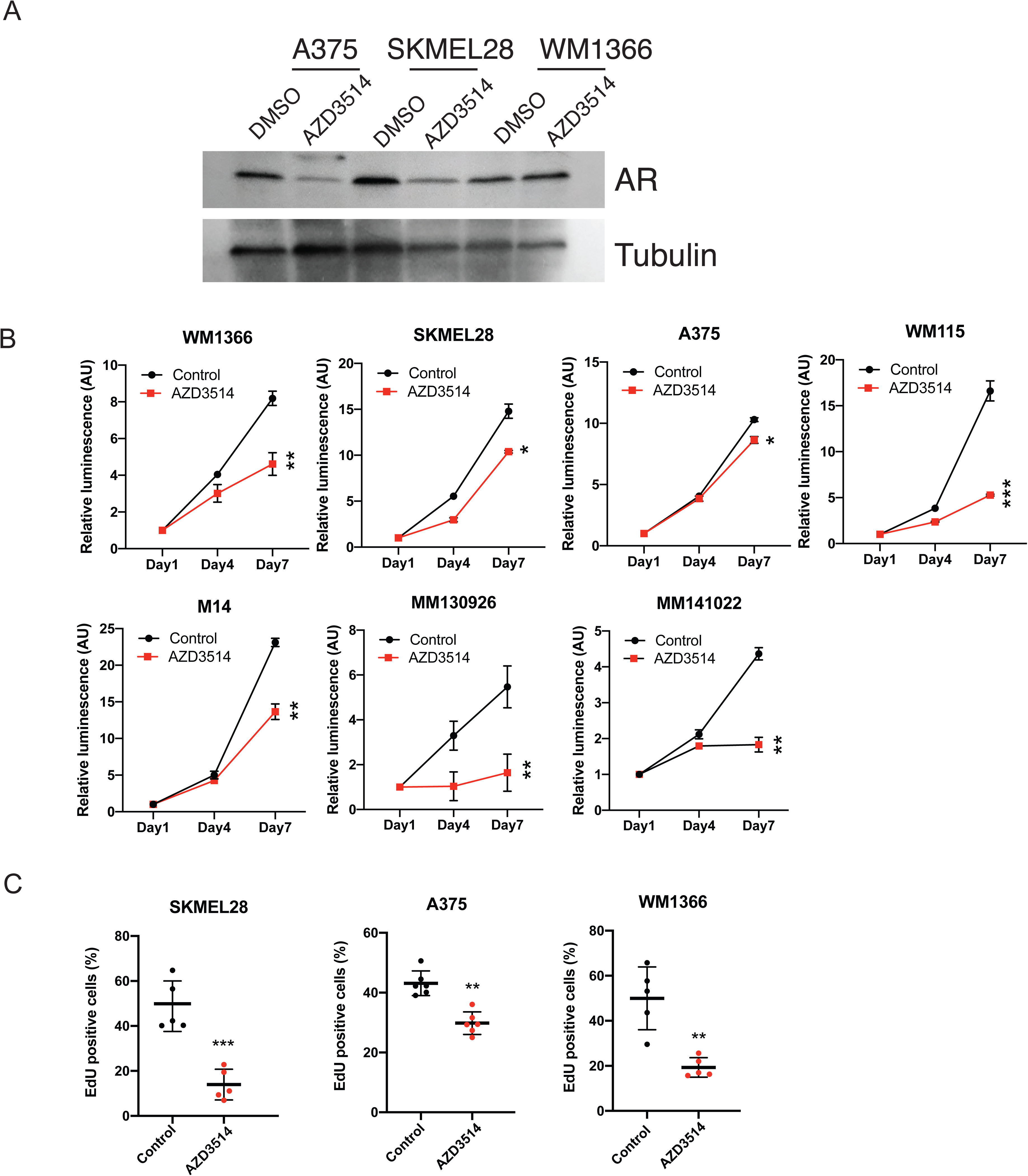
Related to Figure 3B-D. Growth suppressive effects of AR inhibitors on melanoma cells. A) Immunoblot analysis of AR protein expression in the indicated melanoma cell lines treated with AZD3514 (10 µM for 48 hours) versus DMSO control. B) Cell density assays (CellTiter-Glo) of the indicated melanoma cell lines and primary melanoma cells (MM130926, MM141022) treated with AZD3514 (10 µM) versus solvent control (DMSO). Cells were plated on triplicate wells in 96-well dishes followed by cell density / metabolic activity measurements at the indicated days after treatment. Results are presented as luminescence intensity values relative to day 1. C) EdU labelling assays of the indicated melanoma cells treated with AZD3514 (10 µM) versus solvent control (DMSO) at day 5 after treatment. Data are shown as mean ± SD, 1-way ANOVA with Dunnett’s test. n = 3 biological replicates (experiments). *P < 0.05; ** P < 0.01; ***P < 0.005.

**Supplementary Figure 12.**
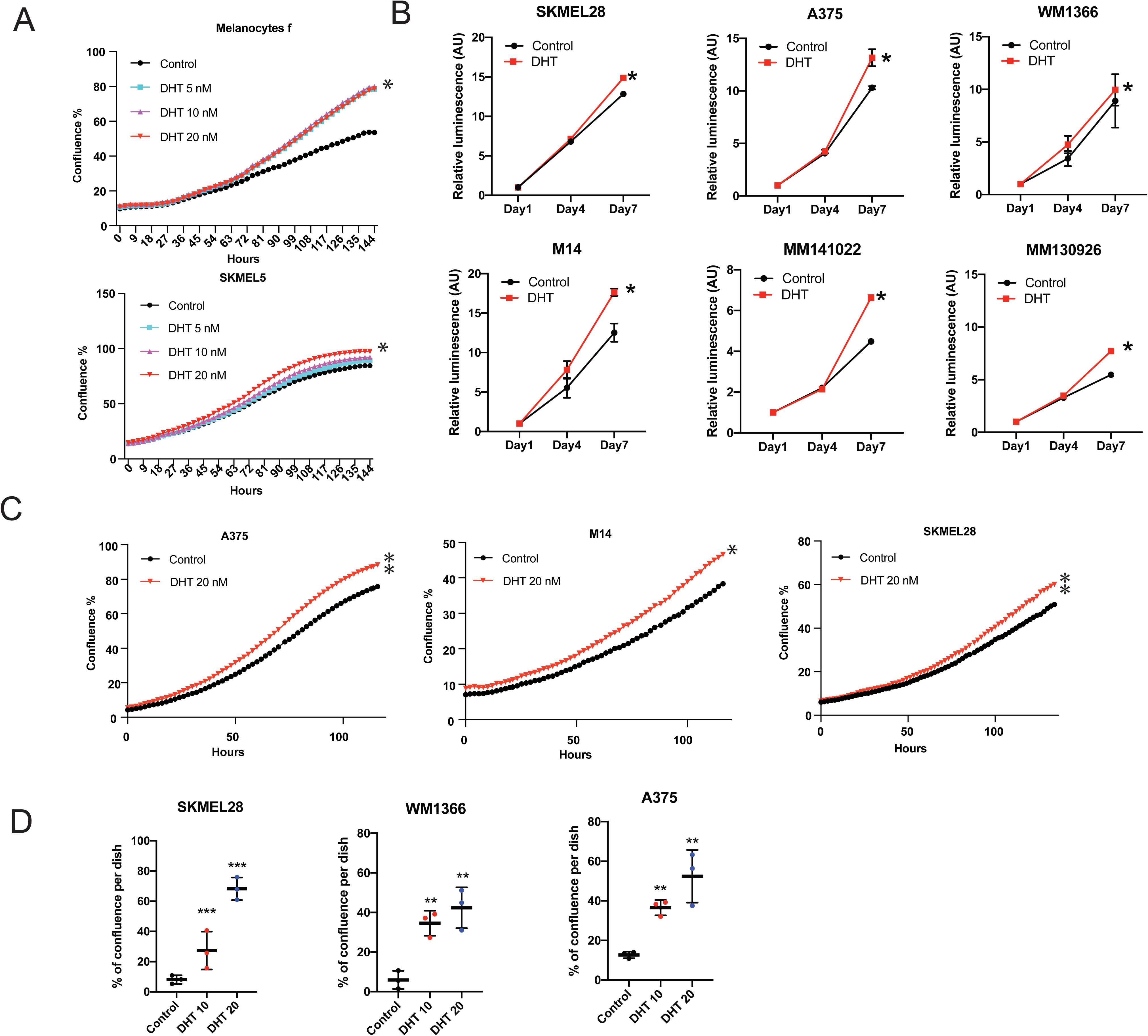
Related to Figure 3E, F. Growth stimulatory effects of dihydrotestosterone (DHT) treatment of melanoma cells. A) Proliferation live-cell imaging assays of the primary melanocytes (strain f) and SKMEL5 melanoma cells treated with the different doses of DHT (5, 10, 20 nM) versus DMSO control by live-cell imaging. Cells cultured in medium with charcoal-treated serum were plated in triplicate wells in 96-well plates followed by cell imaging measurements (IncuCyteTM system, Essen Instruments), capturing 4 images per well every 4 hours for the indicated number of hours. n (number of wells) = 3, Pearson r correlation test. *P < 0.05; B) Cell density assays of the indicated melanoma cell lines and primary melanoma cells (MM130926, MM141022) treated with the AR agonist DHT (20 nM) versus solvent control (DMSO). Cells cultured in medium with charcoal-treated serum were plated on triplicate wells in 96-well dishes followed by cell density / metabolic activity measurements (CellTiter-Glo) at the indicated days after treatment. Results are presented as luminescence intensity values relative to day 1. C) Proliferation live-cell imaging assays of the indicated melanoma cells treated with the DHT (20 nM) versus DMSO control. Assay conditions were as in (A). n (number of wells) = 3, Pearson r correlation test. *P < 0.05; ** P < 0.01. D) Cell density assays of the indicated melanoma cells tested under very sparse condition. Cells were cultured in medium with charcoal-treated serum for 48 hours followed by plating at very low numbers (500 cells in 60 mm-dish) in the same medium plus/minus treatment with DHT (10 and 20 nM) versus solvent control (DMSO) for 7 days. Data are represented as relative cell density as quantified by ImageJ analysis of crystal violet stained dishes. 1-way ANOVA with Dunnett’s test. n = 3 independent experiments. *P < 0.05; ** P < 0.01; ***P < 0.005. All the DHT treatment experiments were carried out in charcoal-stripped medium.

**Supplementary Figure 13.**
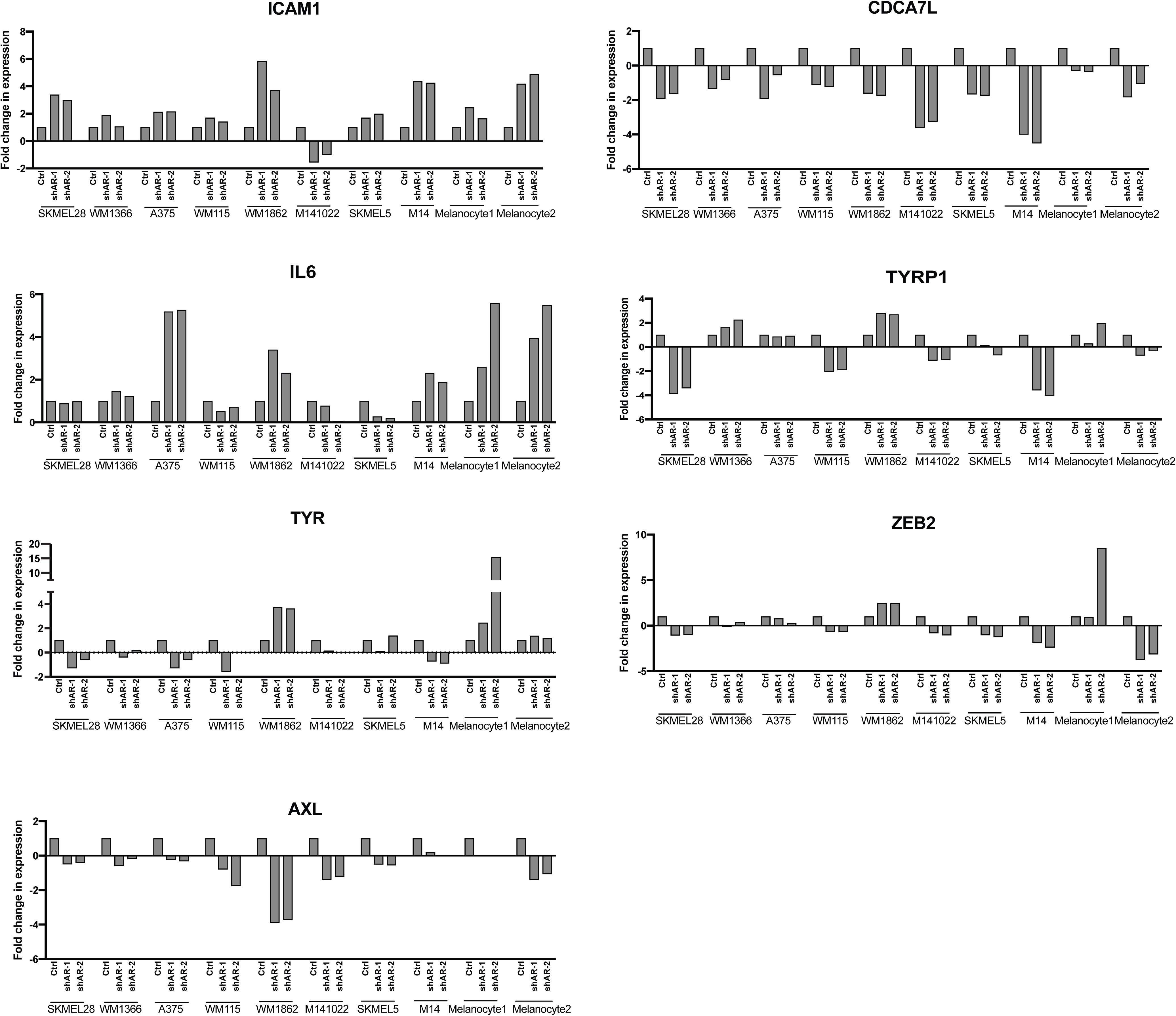
Related to Figure 4B. Impact on melanoma cells gene expression of AR gene silencing. Expression of the indicated genes of interest in a panel of melanoma cell lines, primary melanoma (M141022) and primary melanocytes infected with two AR silencing lentiviruses versus empty vector control, was assessed by RT-qPCR with RPLP0 for normalization. Data, shown as individual plots per cell line, correspond to those shown as heat map in Fig. 4B.

**Supplementary Figure 14.**
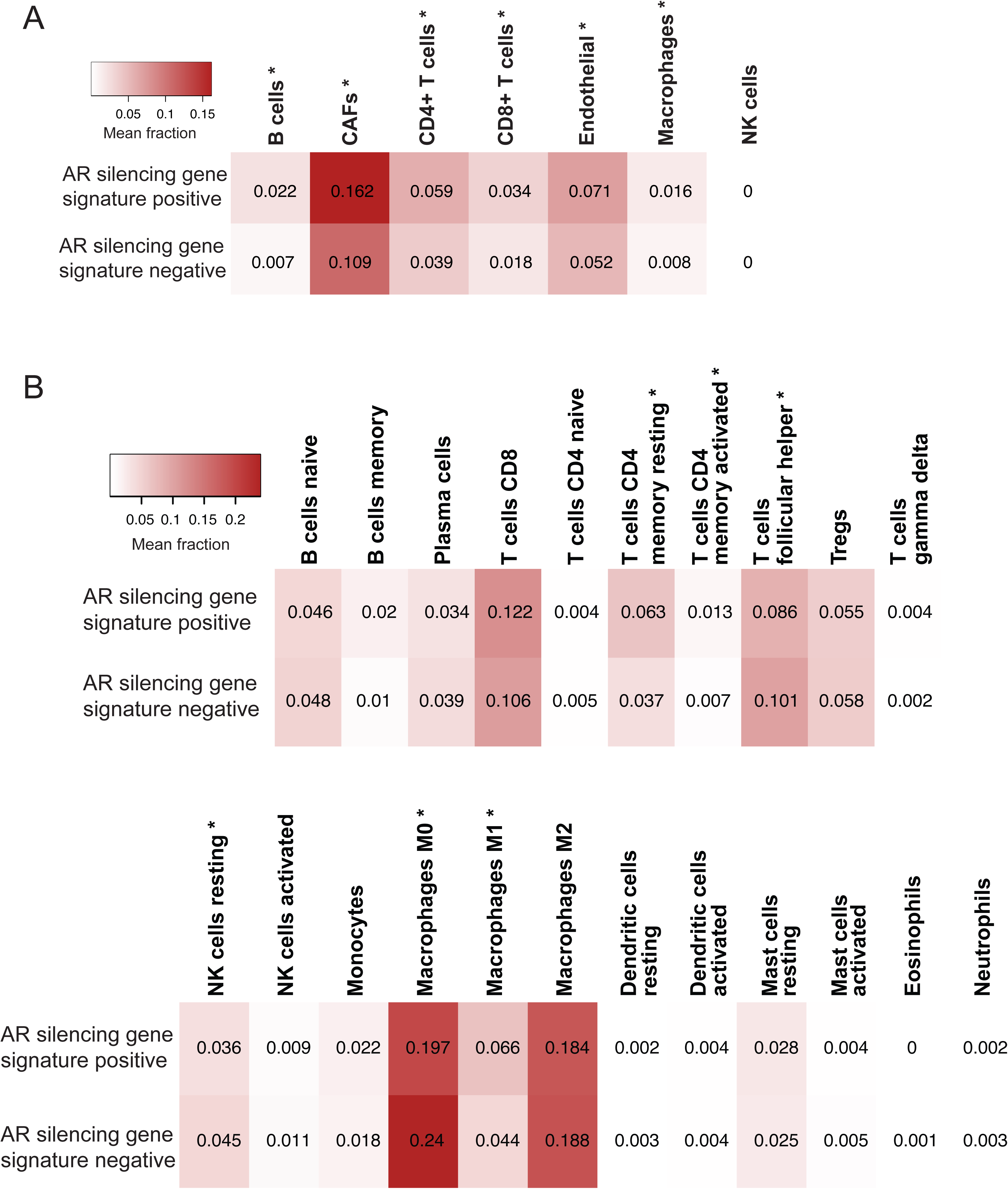
Related to Figure 4E. Prevalence of stromal and immune cells in TCGA-SKCM samples with and without enrichment for the AR silencing gene signature. Heatmaps reporting mean fractions of stromal and immune cell types (columns) for TCGA-SKCM samples with AR silencing signature up or down (rows) obtained using EPIC (A) and CIBERSORTx (B). Red intensity is proportional to the mean cell fraction, which is also reported in each entry. Cell types showing a significantly different prevalence (Wilcoxon rank-sum test, Bonferroni-adjusted p-value < 0.05) between samples with AR silencing signature up or down are highlighted with a “*”.

**Supplementary Figure 15.**
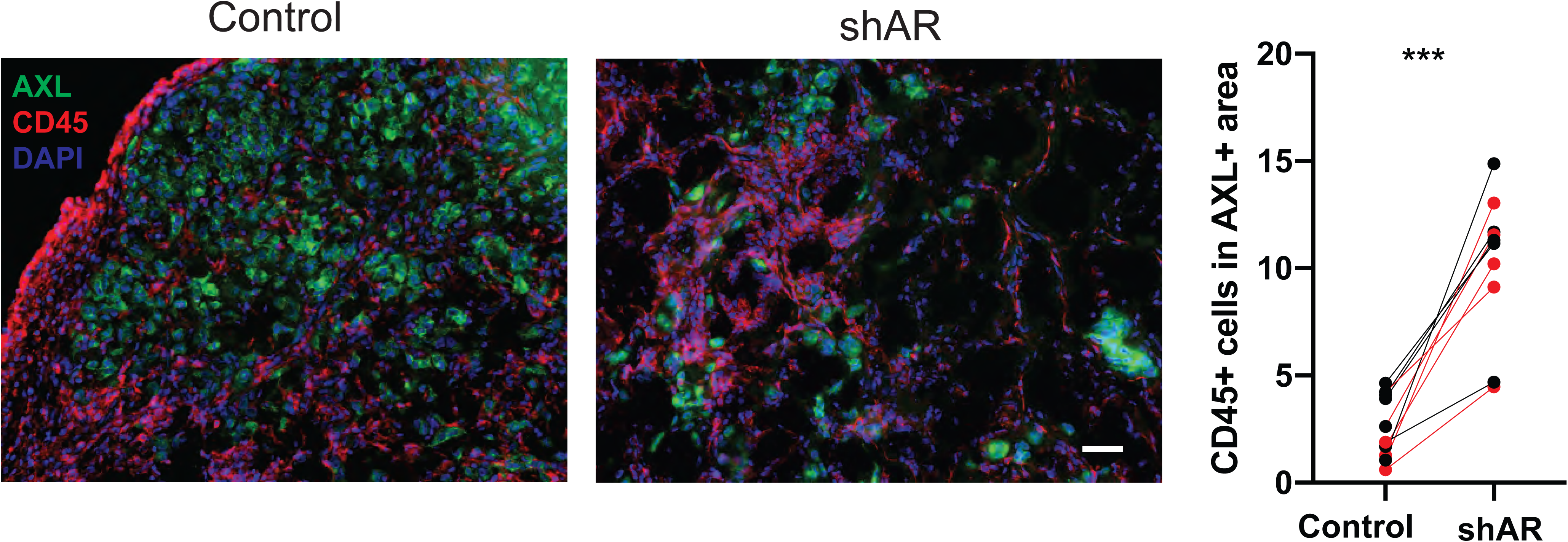
Related to Figure 7. AR silencing inhibits WM1366 melanoma tumorigenesis. Double immunofluorescence analysis of lesions from Figure 7A with antibodies against AXL, for melanoma cells identification and CD45 positive cells. Shown is the quantification together with representative images of CD45 positive cells per AXL positive tumor area, counting in each 5 fields, 5 male mice and 5 female mice, data of male mice in red. Scale bar: 20 µm.

**Supplementary Figure 16.**
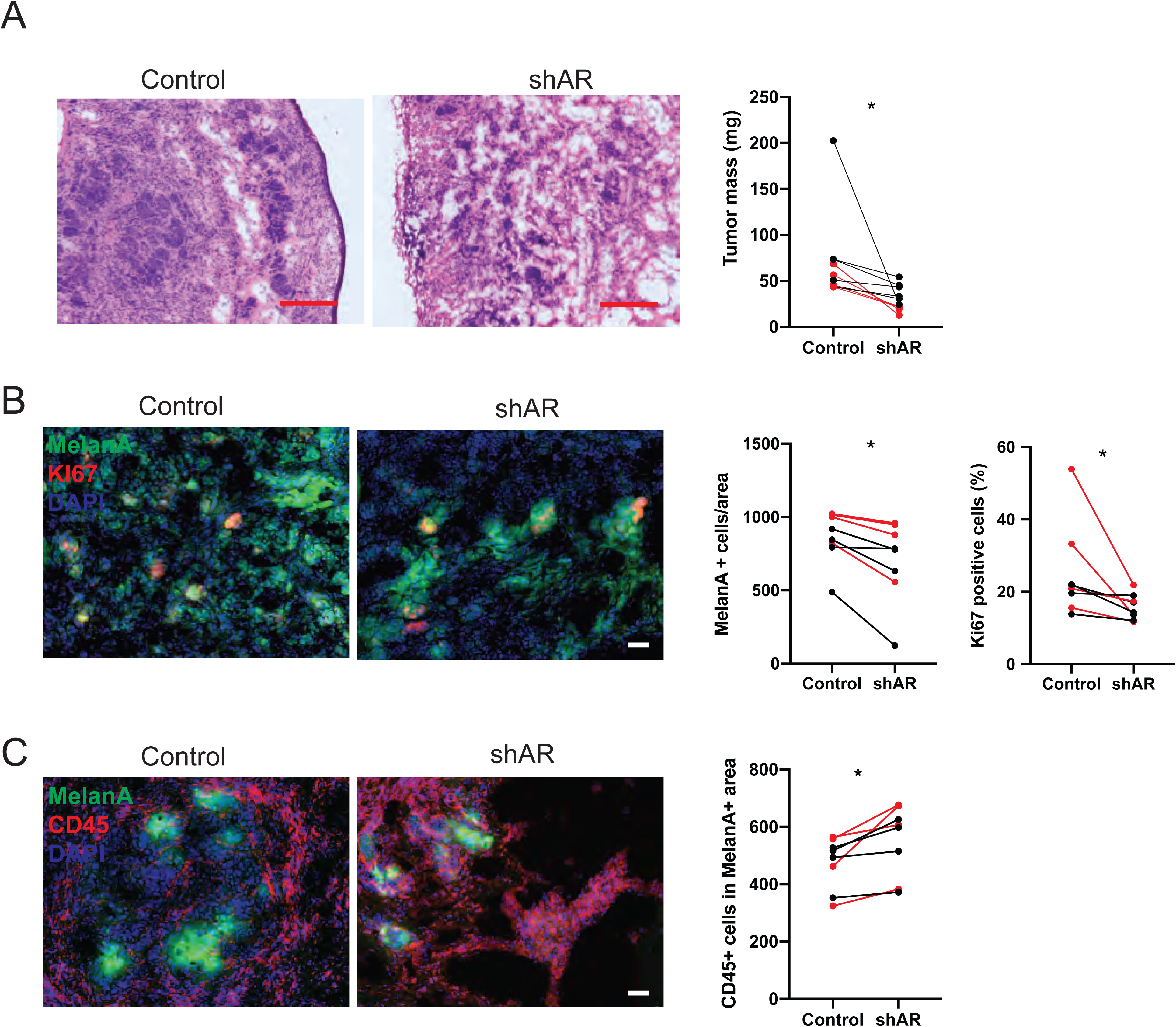
Related to Figure 7. AR silencing inhibits A375 melanoma tumorigenesis. A375 melanoma cells infected with an AR-silencing virus versus vector control were tested by parallel intradermal Matrigel injections into NOD/SCID male and female mice (5 per group, data of male mice in red). Mice were sacrificed 16 days after injection. A) Tumor size, measured by digital caliper (mass = (length x width x height) * π/6) together with representative low magnification H/E images of the retrieved lesions. B) Double immunofluorescence analysis of lesions with antibodies against MelanA (green), for melanoma cells identification, and KI67 (B) positive cells. Shown are representative images of MelanA positive cells stained with antibodies against the other markers, together with relative quantification, (counting in each case >50 cells in 3-5 fields on digitally-retrieved images, using ImageJ software). C) double immunofluorescence analysis of lesions with antibodies against MelanA, CD45 and F4/80, for melanoma cells, hematopoietic cells as well as macrophages identification, respectively. Shown are representative images together with quantification of number of F4/80 positive cells per MelanA positive tumor area, counting in each case 3-4 fields. n (control versus experimental lesions) = 20, two-tailed paired t test, *P < 0.05; ** P < 0.01; ***P < 0.005. Scale bars: 10 µm.

**Supplementary Figure 17.**
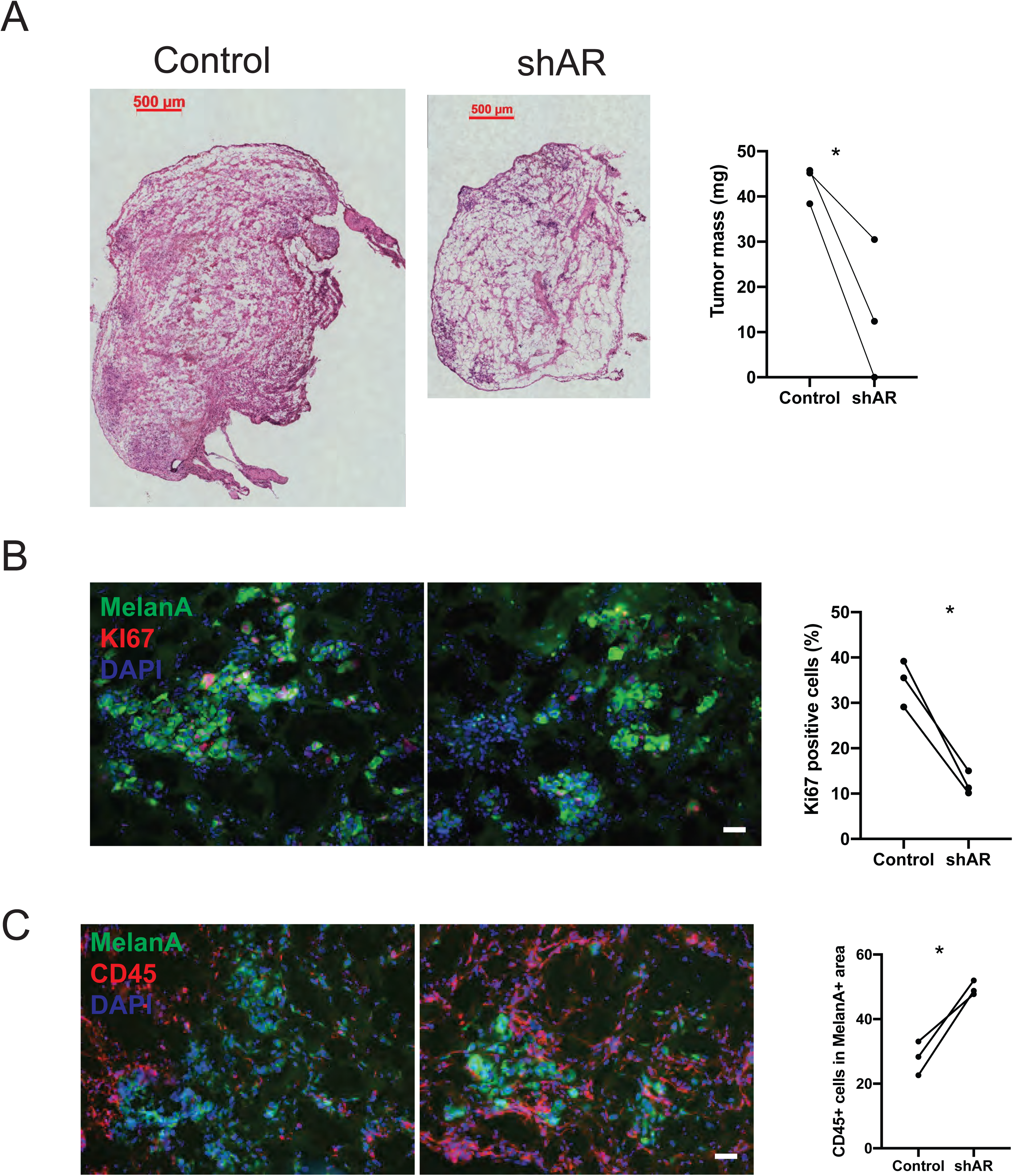
Related to Figure 7. AR silencing inhibits SKMEL28 melanoma tumorigenesis. SKMEL28 melanoma cells infected with an AR-silencing virus versus vector control were tested by parallel intradermal Matrigel injections into NOD/SCID male and female mice (3 per group, male mice). Mice were sacrificed 16 days after injection. A) Tumor size, measured by digital caliper (mass = (length x width x height) * π/6) together with representative low magnification H/E images of the retrieved lesions. Scale bars: 100 µm. B) Double immunofluorescence analysis of lesions with antibodies against MelanA (green), for melanoma cells identification, and KI67 positive cells. Shown are representative images of MelanA positive cells stained with antibodies against KI67, together with relative quantification, (counting in each case >50 cells in 3-5 fields on digitally-retrieved images, using ImageJ software). C) double immunofluorescence analysis of lesions with antibodies against MelanA, CD45 for melanoma cells, hematopoietic cells identification, respectively. Shown are representative images together with quantification of number of CD45 positive cells per MelanA positive tumor area, counting in each case 3-4 fields. n (control versus experimental lesions) = 6, two-tailed paired t test, *P < 0.05. Scale bars: 10 µm.

**Supplementary Figure 18.**
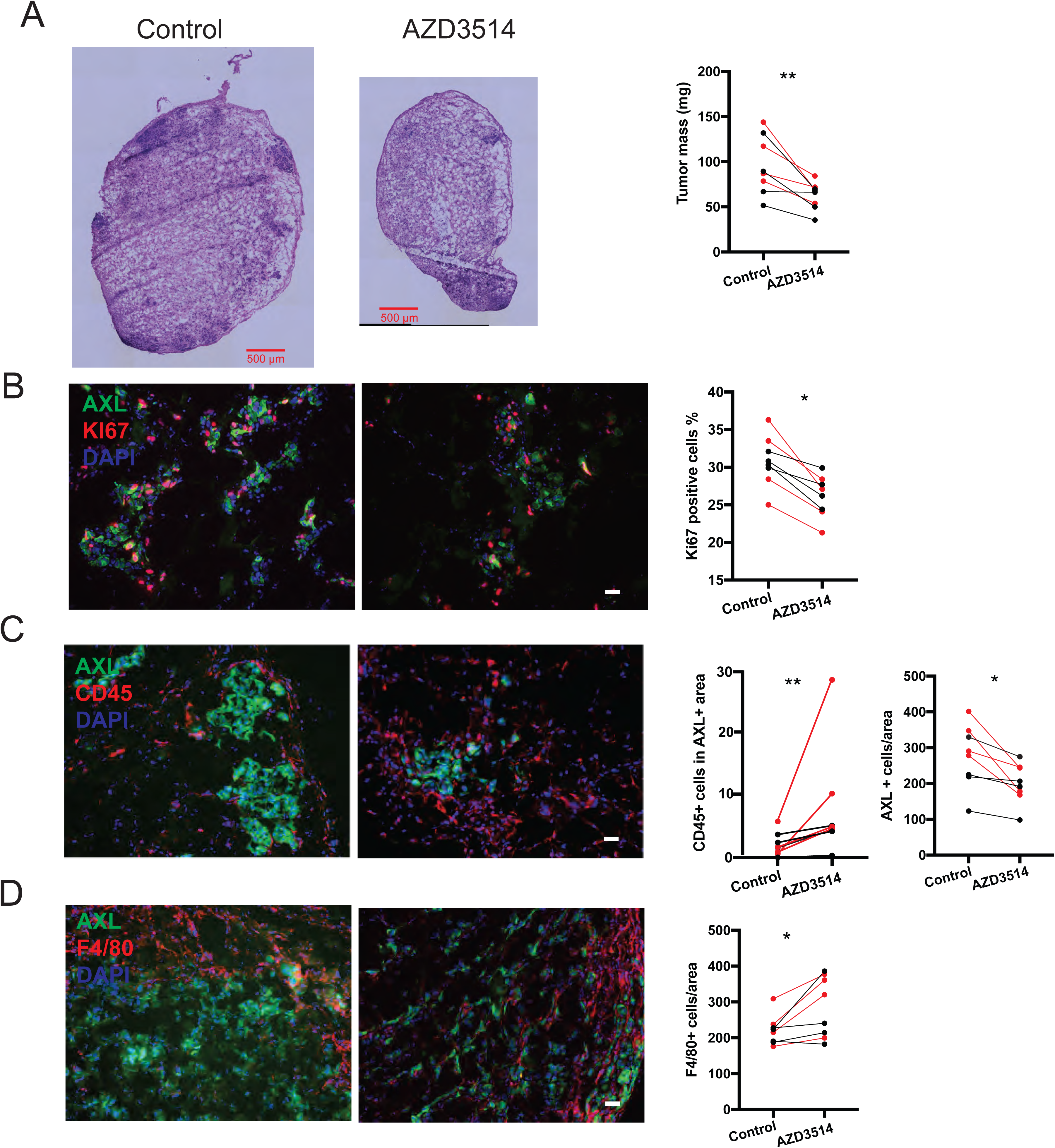
Related to Figure 8. AZD3514 pretreatment inhibits WM1366 melanoma tumorigenesis. WM1366 melanoma cells pretreated with an AR inhibitor AZD3514 versus DMSO control were tested by parallel intradermal Matrigel injections into NOD/SCID male and female mice (4 per group, data of male mice in red). Mice were sacrificed 16 days after injection. A) Tumor size, measured by digital caliper (mass = (length x width x height) * π/6) together with representative low magnification H/E images of the retrieved lesions. B): Double immunofluorescence analysis of lesions with antibodies against AXL (green), for melanoma cells identification, and KI67 positive cells. Shown are representative images of AXL positive cells stained with antibodies against KI67, together with relative quantification, (counting in each case >50 cells in 3-5 fields on digitally-retrieved images, using ImageJ software). C) double immunofluorescence analysis of lesions with antibodies against AXL, CD45 and F4/80, for melanoma cells, hematopoietic cells as well as macrophages identification, respectively. Shown are representative images together with quantification of number of F4/80 positive cells per AXL positive tumor area, counting in each case 3-4 fields. n (control versus experimental lesions) = 16, two-tailed paired t test, *P < 0.05; ** P < 0.01; ***P < 0.005. Scale bars: 10 µm.

**Supplementary Figure 19.**
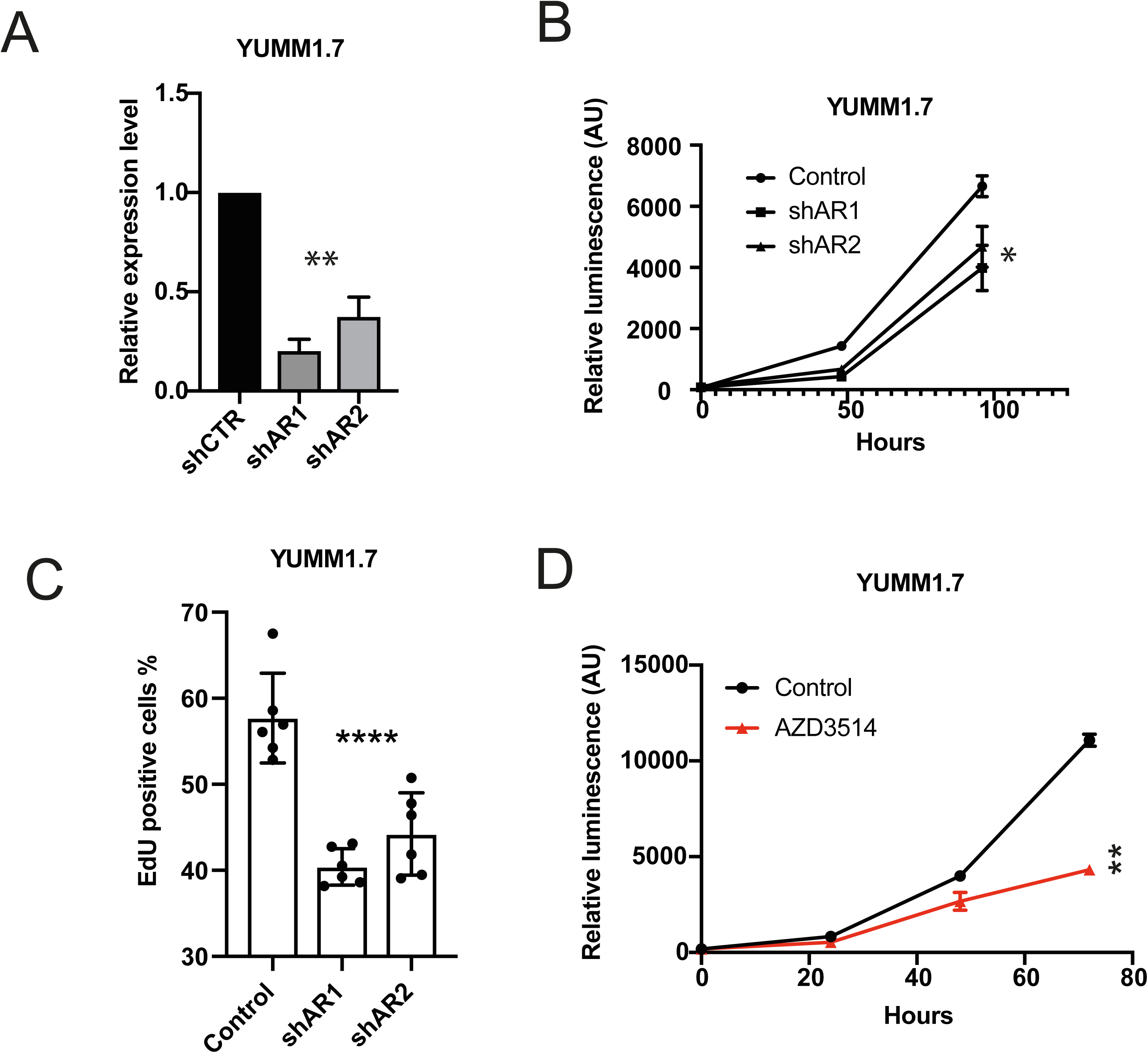
Related to Figure 9. Suppression of mouse melanoma proliferation by AR silencing. A) Level of AR mRNA with shRNA mediated silencing of AR in mouse melanoma cell line YUMM1.7. B) Expression levels of AR mRNA in YUMM1.7 infected with 2 AR silencing lentiviruses versus empty control as assessed by RT-qPCR. Data are shown as mean ± SD, 1-way ANOVA with Dunnett’s test. n = 3 biological replicates (experiments). C) Cell density assays were carried out with YUMM1.7 infected with two AR silencing lentiviruses versus empty vector control. Results are presented as luminescence intensity values relative to day 1. D) YUMM1.7 infected with two AR silencing lentiviruses versus empty vector control were tested by EdU labelling assay 5 days post virus infection. Data are shown as mean ± SD, 1-way ANOVA with Dunnett’s test. n = 3 biological experiments. E) YUMM1.7 were plated on triplicate wells in 96-well dishes followed by CellTiter-Glo metabolic activity measurements at the indicated days after treatment of AZD3514. Results are presented as luminescence intensity values relative to day 1. Data are shown as mean ± SD, 1-way ANOVA with Dunnett’s test. n = 3 biological replicates (experiments). *P < 0.05; ** P < 0.01; ****P < 0.001.

**Table 1. Related to Figure 1. Summary table of patient information of tissue microarray.**

**Table 2. Related to Figure 1. Summary table of a panel of human melanoma cell lines used in this study.** The details of genetic mutations, different clinical histories, AR mRNA levels and growth inhibition/stimulation effects after AR silencing by shAR or AZD3514 (10 µM) and DHT (20 nM) treatment are indicated for each cell type.

**Table 3. Related to Figure 3. List of 155 genes up/down-modulated by AR silencing in three melanoma cell lines (WM1366, SKMEL28 and WM115) from transcriptomic profiling.**

**Table 4. Related to Figure 3. List of gene sets significantly associated with differentially expressed genes in melanoma cells upon AR gene silencing.**

**Table 5. Related to Figure 3. List of perturbagens with concordance with AR signatures including the associated p-values from iLINCS database.**

**Table 6. List of reagents and resources.**

